# At Which Low Amplitude Modulated Frequency Do Infants Best Entrain? A Frequency Tagging Study

**DOI:** 10.1101/2022.12.08.519576

**Authors:** James Ives, Pierre Labendzki, Marta Perapoch Amadó, Emily Greenwood, Narain Viswanathan, Tom Northrop, Sam Wass

## Abstract

Previous infant entrainment research has shown neural entrainment to a wide range of stimuli and amplitude modulated frequencies. However, it is unknown if infants neurally entrain more strongly to some frequencies more than others, and to which low amplitude modulated frequency infants show the strongest entrainment. The current study seeks to address this by testing the neural entrainment of N=23 4–6-month-old infants and N=22 control group adult caregivers while they listened to a range of sinusoidally amplitude modulated beep stimuli at rest (no sound), 2, 4, 6, 8, 10 and 12 Hz. Analysis examined differences across power and phase, regions of interest predetermined by previous literature and by segmented time windows. Results showed that the strongest entrainment was at 2Hz for both adult and infant participants; that there was no significant difference in power and phase, entrainment was occipital temporal and slightly left fronto-central in adults and right fronto-central and left occipito-temporal in infants, leading to some regions of interest used in previous studies being significant in infants and all regions of interest being significant in adults. Segmenting by time window did not show any significant increase or decrease in entrainment over time, but longer time windows showed a stronger entrainment response. In conclusion, it is important to choose appropriate stimulation frequencies when investigating entrainment between stimulation frequencies or across ages; whole head recording is recommended to see the full extent of activation; there is no preference on power vs phase analyses; and longer recordings show stronger effects.

**Author Contribution Statement:** Ives, J., conceptualisation, data collection and curation, formal analysis, methodology, writing – original draft; Labendzki, P., data collection and curation, formal analysis, writing – review & editing; Perapoch Amadó, M., data collection and curation, writing – review & editing; Greenwood, E., data collection and curation, participant recruitment, writing – review & editing; Viswanathan, N., data collection and curation, writing – review & editing; Northrop, T., data collection and curation, participant recruitment, writing – review & editing; Wass, S., conceptualisation, funding acquisition, methodology, project administration, supervision, writing – review & editing.

**Highlights:** 2Hz amplitude modulation stimulation showed the strongest neural entrainment

We discuss power vs phase analyses of infant and adult frequency tagging responses

We illustrate topographic differences in adult and infant neural responses

## 1. At Which Low Amplitude Modulated Frequency Do Infants Best Entrain? A Frequency Tagging Study

Neural activity in mammalian brains during rest and excitation has been shown to take on an oscillatory structure. Oscillations take on the role of pacemaker networks and resonators, each responding to particular firing frequencies (Llinás, 1988). Llinás (1988) describes how these oscillatory networks help specify connectivity during development, help with motor coordination, timing and help with global states, e.g., sleep-wake or attentional state. Buzsáki and Draguhn (2004) demonstrated in the human brain that across five orders of magnitude cortical networks oscillate to help promote processing of incoming sensory information. These neural oscillations help bias input selection, promote long-term consolidation of information and help with time perception.

Person-person and person-environment neural entrainment have been well studied in the adult literature (e.g., Buzsáki and Draguhn, 2004; Glass, 2001; Hoehl, Fairhurst & Schirmer, 2021; Thut, Schyns & Gross, 2011) and to some extent infant literature (Wass et al., 2020; Wass, Perapoch Amadó & Ives, 2022). Neural entrainment is measured as matching periods of oscillatory activity, measured either concurrently or simultaneously across a dyad, as a result of a shared stimulus or social interaction, producing a bidirectional influence between the dyad (Wass et al., 2020).

Entrainment studies in adults have suggested that entrainment helps attention selection (Besle et al., 2011; Lakatos et al., 2008; Thut, Schyns & Gross, 2011; Ward, 2003), memory encoding (Ward, 2003), memory recall (Hickey et al., 2020), sensory selection (Schroeder & Lakatos, 2009), auditory perception (Kayser et al., 2015; Rimmele et al., 2018), music processing (Doelling & Poeppel, 2015), speech processing (Ding et al., 2015; Giraud & Poeppel, 2012; Hyafil et al., 2015), interperson conversational features including speech pitch, intensity voice quality and speaking rate (Levitab & Hirschberg, 2011) audio-visual processing (Schroeder et al., 2008), synchrony during face-to-face communications (Jiang et al., 2012), visual-olfactory processing (Rekow et al., 2022), prediction of future actions (Kayhan et al., 2022), coordination of movement and auditory rhythm (Varlet, Williams & Keller, 2018) and event timing (Kösem, Gramfort & van Wassenhove, 2014) (for further review see Kabdebon et al. (2022).

Many behaviours are thought to evoke neural entrainment including conversations (Pérez, Carreiras & Duñabeitia, 2017) that feature both infant directed and adult directed speech (Kalashnikova et al., 2018), singing (Weinstein et al., 2016) listening to audiobooks (Koskinen & Seppä, 2014), watching movies (Lankinen et al., 2014) and watching cartoons (Jessen et al., 2019). Entrainment is also thought to be coupled across many physiological markers, including mother-infant synchronisation of heart rhythms (Feldman, 2007), neural tracking of pain (Guo et al., 2020), thermal-cardiac entrainment (Mannix et al., 1997), infant-adult synchronisation of pupil dilations (Fawcett et al., 2017) and synchronisation of respiration to rocking (Sammon & Darnell, 1994).

For further review of entrainment in early social interaction, and oscillatory entrainment to early social and physical environments please see Wass et al., (2020) and Wass, Perapoch Amadó and Ives (2022).

### 1.1. Developmental Auditory Capabilities

The ongoing auditory scene from our surrounding environment is a constant barrage of overlapping spectral and temporal information that arrives at our ears as a single stream, but which humans separate into distinct sources during auditory processing in the brain. Näätänen et al., (2001) describe how adults use “sensory intelligence” to untangle this information to produce an auditory scene in the auditory cortex. Young infants, including newborns (McAdams, 1997; Winkler et al., 2003) and 7–15-week-olds (Demany, 1982), have been shown to separate concurrent audio streams of information, suggesting that some elements of how complex auditory sounds are processed seem likely to be present at birth. After grouping and organising the auditory input, humans have then been shown to track and apply hierarchies to the information. For example, this is commonly seen in the cortical tracking of speech structures in adults (Ding et al., 2016) and infants (Choi et al., 2020; Leong & Goswami, 2015). This is also seen across sensory modalities spanning very low frequencies (< 1Hz), including across minutes, hours and days (for a review see Wass, Perapoch Amadó & Ives, 2022).

Infants, even newborns, have been shown to be born with a range of complex auditory processing capabilities that allow them to segregate sounds and organise their auditory world. For example, infants can detect changes in pitch (Alho, 1989), prefer their native maternal language (Moon, Cooper & Fifer, 1993), and recognise and prefer their mother’s voice (De Casper & Fifer, 1980). Machine learning has been shown to reliably classify the neural response of 8-week-olds to both rhythmic and non-rhythmic speech (Gibbon et al., 2021). Using this segregated information, newborns have been shown to model acoustic regularity by learning to group repeating tones out of random patterns (Stefanics et al., 2007), and model acoustic regularity and show a neural response to violations in expected auditory sequences (Carral et al., 2005).

At a more granular level, infants have been shown to employ statistical learning as patterns to perceive acoustically relevant information. Saffran et al., (1999) demonstrated the first evidence of infant statistical learning of tone sequences in both infants and adults. Further studies have demonstrated that infants under the age of 12 months can use statistical learning to pick out words in regular computerised speech (Choi et al., 2020; Fló et al., 2022; Kabdebon et al., 2015), while cortical tracking studies have demonstrated that infants track sung speech (Attaheri et al., 2022), cartoons (Jessen et al., 2019) and infant directed and adult directed speech (Kalashnikova et al., 2018).

### 1.2. Infant vs Adult Neural Entrainment Capabilities

Auditory steady state responses (ASSRs) are cortical responses to fast isochronously repeating stimuli, which are often used to test hearing capabilities in newborns and young infants. ASSRs are a valuable technique as they can be administered to adults and infants whether they are awake or asleep (e.g., Cohen et al., 1991; Jerger et al., 1986; Wang et al., 2022). Using ASSRs, infants have been shown to cortically track simple stimuli from a very early age (e.g., Lorenzini et al., 2022). Daneshvarfard et al., (2019) demonstrated that preterm infants even at an average gestational age of 31.48 weeks exhibited a classic ASSR. While Niepel et al., (2020) showed using foetal MEG that infants as young as 30 weeks gestational age exhibit ASSRs *in utero*, suggesting that even with limited capacity to interact with the outside environment foetal brains can track and entrain to rhythmic stimuli. However, while there is evidence that infants can entrain to auditory stimuli as a method of parsing auditory information in a more efficient manner, there is clear evidence that adult and infant auditory processing is not the same, and entrainment studies using adults cannot be used as a proxy for infant auditory entrainment (for a review of dissimilarities between adult and infant auditory processing please see Saffran, Werker & Werner, 2006).

One expected contributor, especially during early infancy, is the maturation occurring in infant auditory cortices and neural networks (e.g., Adibpour et al., 2020). This, in turn, impacts infant’s ability to neurally synchronise with their environment and the people around them (Shafer, Yu & Wagner, 2015; Uhlhaas et al., 2010). As an example, the original ASSR study in adults by Galambos et al., (1981) used as a technique to test auditory pathways, showed a distinctive response at 40Hz and was quickly taken up globally (Kuwada et al., 1986; Rees et al., 1986; Rickards and Clark, 1984; D. Stapells et al., 1984). However, there is not such a strong response in infants at higher frequencies including at 40Hz (Maurizi et al., 1990; Picton, 2003). Instead, this response grows as children mature into adults (Rojas et al., 2006) due to the protracted maturation of the auditory cortex (e.g. Adibpour, 2020), where the 40Hz ASSR has been reported to be centred in response to clicks, noise bursts (Hari, Hämäläinen, & Joutsiniemi, S., 1989) and sinusoidal tones (Herdman et al., 2002).

Infants have been shown to entrain at a range of lower amplitude modulation frequencies (see section 1.4) and have been shown to mentally construct meter in scenarios where there is an ambiguous meter (Cirelli et al., 2016). The ability to entrain to these low frequency rhythms may be a direct result of low amplitude modulated stimuli targeted at infants from adult caregivers, which has been shown to consistently change to match an infant’s abilities across many languages (Narayan & McDermott, 2016).

### 1.3. Why are slow amplitude modulated rhythms important?

Sounds include two key oscillatory components, a carrier frequency, which is often referred to as pitch, and an amplitude modulated frequency, often referred to as the rate, tempo or rhythm of a repeating sound. Carrier frequencies must be above 20Hz to fall into the audible range of humans, this overlaps with the amplitude modulated frequencies which start at rates below 0.1Hz and can reach into the hundreds of Hz. The point at which amplitude modulated frequencies become carrier frequencies is the point at which peaks, and troughs of the oscillations are no longer individually discernible from neighbouring oscillations.

Slow amplitude modulated rhythms are prevalent in natural sounds, which may carry critical information (Singh et al., 2003). They are a key component of human social interactions such as speech and have consistently been shown to encode temporal cues that help people forward predict social information during social interactions (Ahissar et al., 2001; Alaerts et al., 2008; Aiken & Picton, 2008; Bertoncini et al., 2011; Henry, Herrmann and Grahn, 2017; Wang et al., 2011). Greenberg et al., (2003) demonstrated that properties of speech under 5Hz generally represent a lower branch of modulated speech, used for heavily stressed syllables, while speech properties with higher frequencies between 6-20Hz generally reflect unstressed syllables. Goswami and Leong, (2013) investigated the characteristics of speech and suggested that the stress, syllable and part of the phoneme presentation rate were under 20Hz.

Similarly, when phase locking value (PLV) of neural signal to white noise between 4-128Hz was calculated by Liegeois-Chauval et al., (2004) the auditory cortices were shown to have the strongest response to the lowest amplitude modulated frequencies between 4-16Hz, matching the range crucial for speech intelligibility.

While in the process of decoding the complex stimuli around an infant that help infants to develop the skilled task of contingent social interactions, it seems especially important that infants are able to neurally entrain to a range of low amplitude modulated signals. This is important in speech, as shown above, but also in the effect of a wider range low amplitude and ultra-low amplitude (< 1Hz) modulated oscillators including respiratory, arousal, sleep-wake and hormone cycles (Feldman, 2006; Feldman, 2007) on infant-caregiver synchrony. In the words of Feldman (2006) “the organization of physiological oscillators appears to lay the foundation for the infant’s capacity to partake in a temporally matched social dialogue”.

### 1.4. Research of Stimuli Frequency

Given the importance of low amplitude modulated auditory frequencies to infant development, it is important to ask to which frequencies do infants best entrain? While there have been some infant-adult or infant studies, the question of entrainment to auditory stimuli has received much more attention in the adult literature.

Many MEG and EEG studies have examined these questions from the perspective of hearing acuity or auditory perception: Luo et al., (2006) investigated phase vs power neural tracking between 0.3-8Hz; Nozaradan, Peretz and Mouraux, (2012), studied neural entrainment to beat and meter in musical rhythms between 0.426-5Hz; Rees, Green and Kay, (1986) looked at the effects of modulation depth on ASSR responses between 0.4-400Hz; Wang et al., (2012) studied sensitivity to spectral bandwidth in speech processing between 1.5-31.5Hz; Picton et al., (1987) examined the effects of amplitude modulation vs frequency modulation in the context of modulation depth between 2-12.7Hz; Peelle, Gross and David, (2013) examined neural responses to speech amplitude envelope between 4-7Hz; Millman et al., (2010) investigated the spatiotemporal structure of the auditory steady state response (ASSR) at 4, 8 and 12 Hz; Henry, Herrmann and Grahn (2017) examined responses to tone duration, onset/offset duration and input patterns between 5-40Hz; Roß, Borgmann and Draganova, (2000) investigated magnitude of neural response between 10-98Hz; Kuwada, Batra and Maher, (1986) looked at the impact of steady state responses on normally hearing vs hearing-impaired participants between 25-350Hz, with the largest steady state responses found between 25-50Hz.

There have been a few studies that have examined adults and infants including: Stapells et al., (1988) who tested 3 week to 28-month-old infants with a variety of steady state responses between 9-59Hz in 5Hz steps and showed no consistent neural peak. Levi, Folsom and Dobie, (1993) presented adults and 1 month old infants with stimuli amplitude modulated between 10-80Hz.

Adults showed an increased magnitude of response at 40Hz which then dropped for later frequencies, while infants showed an increased linear response between 10-80Hz. Aoyagi et al., (1994) studied auditory sweeps of frequencies rapidly presented between 20-200Hz in 20Hz steps to gauge hearing acuity as a function of age in participants between 4 months to 15 years and a cohort of adults showing that the optimal steady state response was detected at 80Hz for young infants (2-4 years old). Riquelme et al., (2006) tested 149 newborns with a range of stimuli including carrier frequencies of 500, 1KHz, 2KHz and 4KHz that were amplitude modulated between 25-98Hz to test for the presence and magnitude of steady state responses. Riquelme et al., (2006) also tested the impact of intensity of these stimuli on the infants with stimuli volumes between 20-70dB. They demonstrated that there were steady state responses between 41-88Hz across all carrier frequencies. Pethe et al., (2004; see also Savio et al., 2004) investigated the impact of age on two frequencies, 40 and 80Hz, with infants between 2 months and 14 years old. Pethe et al., (2004) found that the optimal steady state response changed from 80Hz to 40Hz for children aged 13 and above, matching adult responses. Finally, Rickards et al., (1994) investigated the impact of modulation depth on 337 sleeping newborns between 60-100Hz, finding steady state responses at all frequencies tested to varying degrees.

The above research demonstrates that there have been investigations into the impact of stimuli frequency both in adult and infant studies. However, there are two issues with the above studies in relation to the current question. First, the majority of these studies, especially the infant studies, tested frequencies much higher than the low amplitude modulation rate important in speech and speech perception. Second, most of these studies have investigated a range of frequencies for the primary purpose of hearing acuity tests, which use a small number of cycles for each stimulation frequency. While there may be evidence of entrainment at these frequencies it is difficult to measure the extent to which these are the result of endogenous oscillators entraining to the stimuli or exogenously generated stimulus responses (Haegens & Golumbic, 2018; Wass, Perapoch Amado & Ives, 2021). It is expected that there will be a stimulus response to any stimuli in the environment that has a high enough intensity or salience, but this does not show the neural tracking of these stimuli after initial habituation, i.e. after the reduction of high magnitude exogenous stimulus responses, driven by a high level of attention to a new stimulus.

### 1.5. Measuring Infant Neural Entrainment with Frequency Tagging

#### 1.5.1. Frequency Tagging

There are multiple ways to investigate the characteristics of infant neural entrainment, one popular method is frequency tagging. Fundamentally, frequency tagging assumes that the neural response to a rhythmic stimulus will be at the same frequency as the stimulation frequency. This allows researchers to “tag” a particular response in a specific brain region. The enticing aspect of frequency tagging is that the stimulus characteristics can be manipulated to test perceptual capabilities (e.g. complex rhythm detection, Nozaradan et al., 2017), test high and low order comprehension (e.g. hierarchy of linguistic components in speech, Lo et al., 2022) and can be completed with multiple stimulus types in parallel (e.g. social vs non-social stimuli, Vettori et al., 2020). Frequency tagging requires minimal effort from participants, and in the case of auditory frequency tagging can be completed while participants sleep, as sleep has been shown to dull but not remove steady state responses (e.g. Cohen et al., 1991; Jerger et al., 1986; Wang et al., 2022).

#### 1.5.2. Stimulus Characteristics

Stimulus characteristics have been explored in previous literature. Zhou et al., (2016) have investigated the interpretation of expected responses to single stimuli vs repeated periodic stimuli, single stimulus duration and effects of fast onset/offset stimuli vs sinusoidal amplitude modulated stimuli, as well as expected harmonic responses using stimuli with varying characteristics. While Kabdebon et al., 2022 have suggested considerations for the stimulation rate in frequency tagging studies, e.g. the stimulation frequency should bear in mind the cognitive response time of the region of interest being targeted; frequencies relating to dominant neural frequencies (6-9Hz in infants, 9- 12Hz in adults) should be avoided if possible due to the higher resting power at these frequencies; and that there should be at least 4-8 frequency bins between stimulation frequencies if multiple stimuli are being tested. However, while previous research has suggested frequencies to avoid to stop the response clashing with dominant neural frequencies, to the best of our knowledge, there have been no studies that advise on which frequencies show the strongest entrainment.

#### 1.5.3. Analysis method

A tagged response can be seen both in the power and phase domains (for a review see Kabdebon et al., 2022). However, it is not clear from this review, which method of analysis should be preferred, whether both are needed or if the methods could be used interchangeably. Zoefel, ten Oever and Sack, (2018) describe one issue with using a Fourier transform to measure neural frequency tagged response. Neural mechanisms such as phase resetting of neural signals to better entrain to an ongoing stimulus violate the stationarity of the signal, so when compared to a sinusoid, e.g. during a fast Fourier transform, the power of the signal will be lowered. This may also represent an issue for phase analyses if the phase resetting produces phase angles that are opposed to one another (e.g. 180 degrees apart), as this may make the amplitude of the phase measure cancel itself out, e.g. during a biopolar or tripolar phase angle response.

One potential solution to this is to use entropy of phase angle. Entropy measures the variance in responses rather than a cumulative phase angle and so responses with bipolar or tripolar opposites would not be cancelled out. Entropy of phase angle has been used previously by Notbohm, Kurths and Herrmann, (2016) when measuring the impact of stimulus intensity on neural entrainment.

### 1.6. Auditory Frequency Tagging Regions of interest

Previous studies have focused on particular regions of interest when conducting auditory frequency tagging. Many of these relate to auditory centres of the brain, especially when the objective of the study is to use auditory frequency tagging as a measure of auditory perception. These include Cz only (e.g. Aoyagi et al., 1994; Cohen, Rickards and Clark, 1991; Jerger et al., 1986; Mühler, Rahne and Verheya, 2013; Pethe et al., 2004; Stappels et al., 1988); Forehead/FPz only (e.g. Alaerts et al., 2009; Levi, Folsom and Dobie, 1993; Maurizi et al., 1990; Rickards et al., 1994; Riquelme et al., 2006; Savio et al., 2001); vertex area (e.g. Choi et al., 2020; Rojas et al., 2006); surrounding the zenith line from anterior to posterior (e.g. Daneshvarfard et al., 2019; Ramos-Escobar et al., 2021); and whole head recordings (e.g. Attaheri et al., 2022, Choi et al., 2020, Cirelli et al., 2016; Herdman et al., 2002; Kabdebon et al., 2015).

Choosing a region of interest is often part of a hypothesis driven approach. Investigating commonly known regions of activity (e.g. electrodes corresponding to A1, Wernicke’s area, Broca’s area or the perisylvian region), using previous research highlighting topographic activity or reducing unnecessary topographic artefact are seen as methods to reduce fishing for spurious results.

However, there is also the possibility to miss potentially interesting regional activation when investigating a topic where there is limited knowledge, especially as there are developmental changes in the topographic activity due to broader developmental change.

### 1.7. Current Study Aims

The current study aims to fill the gap in this research area by testing infants and adults with a range of low amplitude modulated frequencies to investigate their neural entrainment responses and determine if there is a preference for any particular frequency. It is expected that there will be an increased response at the dominant frequencies for infants (6-9Hz) and adults (9-12Hz), which will be seen in both power and phase analyses. This is thought to be true because these dominant frequencies can be flexible to help track oscillations in the environment (for a review see Zoefel, ten Oever and Sack, 2018), but increasing the spectral distance between two oscillators increases the intensity of the stimuli to bring two oscillators in alignment, which is the fundamental concept of the Arnold’s tongue (Notbohm, Kurths, Herrmann, 2016; Zoefel, ten Oever & Sack, 2018).

We anticipate that there will be differences in the power and phase response; responses over the time course of the stimulation; and with regional differences across the scalp. Using the frequency that shows the strongest response for each participant group, the current study will therefore investigate differences in power vs phase, temporal and spatial responses. We realise that this investigation will not be generalisable to all studies due to the broad range of stimulus types, infant participant ages and experimental settings. This means that we cannot provide authoritative guidance on the best approach to all research scenarios. However, we expect to learn lessons that will be valuable to the infant development, neural entrainment and frequency tagging communities when planning similar studies.

## 2. Method

### 2.1. Participants

26 adult-infant dyad participants were recruited as part of a wider longitudinal project. Of these, 7 infant datasets were rejected due to poor data quality, 6 adult datasets were rejected due to technical fault or poor data quality. The study included 7 conditions: no sound (rest), 2Hz, 4Hz, 6Hz, 8Hz, 10Hz and 12Hz. After preprocessing of EEG data, infants contributed 6, 8, 11, 10, 12, 9 and 9 participants to these passive audio conditions respectively, while adults contributed: 6, 7, 10, 12, 18, 6 and 10 participants to the same conditions, a table detailing which participants completed which conditions has been placed in the supplementary materials. Conditions were presented in a random order, testing was stopped if the child fussed or woke up, contributing to the uneven sample sizes. Participant information has been included in the supplementary materials.

Mean adult participant age was 35.53 years (standard error, 0.698), mean infant age 5.39 months (range 4.46-6.36 months, standard error 0.099). All adult participants were female, infant participants included 16 male, 10 female. Infants had an average gestational age of 40.51 weeks.

The University of East London ethics committee approved the study. All adult participants provided informed consent for both themselves and their children according to the Declaration of Helsinki. All participants were offered a £10 shopping voucher as a monetary reward for their time. Travel and food expenses were also covered for those that requested them.

The current experiment was part of a longitudinal research project, which tests infant-caregiver dyads at 5-, 10-, 15- and 36-month timepoints. As part of the testing protocol a lab session is run split into three sessions. The data for this study was one of the sessions, the order of which varied depending on when the infants slept during the day.

### 2.2. Stimuli

Audio stimuli were generated in MATLAB 2021b, consisting of pure sinusoidal tones using a carrier frequency of 1000Hz and sinusoidal amplitude modulation at either 2, 4, 6, 8, 10 or 12Hz saved with a sampling frequency of 48KHz. Audio stimuli had a minimum amplitude of -1 and a maximum amplitude of 1, which was then controlled to the target volume of 65dB using on speaker controls and a sonometer. Each audio file was created with 240 seconds of continuous audio.

### 2.3. Procedure

Data collection was conducted when an infant fell asleep during a lab testing session. The experimental environment changed depending on how the infant was comfortable sleeping. Infants slept in their mothers’ arms, in a moses basket, on a sofa, baby sling/carrier or in their pram. The room was kept quiet and in the majority of cases the lights were dimmed to help the child sleep. No auditory sleep aids were permitted.

Creative SBS 250 speakers were placed close to the participants and calibrated to 65dB at the participants’ ear using a RS PRO RS-95 sonometer, chosen to be at the average volume of standard speech (e.g. Olsen, 1998).

Adult participants were also instructed to listen to the passive audio task and were permitted to sleep. Participants were asked not to do any rhythmic motions including: chewing, talking, fidgeting, waving, bouncing etc, and asked not to eat or use any electronic devices. Participants were told that if they wanted to stop for any reason or if the infant woke up then the procedure would be stopped.

### 2.4. EEG Data Acquisition

EEG signals were obtained using a dual BioSemi (Amsterdam, NL) ActiveTwo system configured for 64 channel recording from both participants simultaneously. Participants wore size appropriate 64 channel Electro-Cap International (Ohio, US) caps with a 10-20 electrode montage. Common Mode Sense and Driven Right Leg electrodes between Pz and POz were used as the active reference. EEG signals were recorded at 512Hz with no online filtering using ActiView data acquisition software (version 7.07; BioSemi). Signa Gel conductive electrode gel from Parker Laboratories BV (Almelo, NL) was used to bridge the connection between the electrodes and the participant’s scalp.

### 2.5. EEG Artifact Rejection and Preprocessing

A modified version of Marriott-Haresign et al.’s, (2021) automated EEG artifact rejection and pre-processing pipeline was implemented. First the data were high pass filtered at 1Hz using the EEGLAB (Delorme & Makeig, 2004) function *pop_eegfiltnew.m* (Widmann, 2008; default settings, filter order 3380). Second, line noise was removed using a notch filter with *pop_cleanline.m* (Mullen, 2012). Third, a low pass filter was applied at 25Hz with the same settings as the highpass filter.

Fourth, the data were referenced to a robust average, calculated by first temporarily removing noisy channels using *clean_channels.m* (Kothe, 2014; default settings) and averaging the remaining channels. Fifth, after averaging all channels using the robust average, channels that were subsequently still noisy were rejected using *clean_lines.m* with a correlation threshold of 0.7 and a noise threshold of 4. Subsequently, all channels that were not rejected were put through *eBridge.m* (Alschuler et al., 2014; default settings) to identify bridged electrodes. All electrodes that were identified as noisy or bridged were interpolated using the spherical method of *eeg_interp.m* (Delorme, 2006). Sixth, using a sliding 1 second window without overlap, epochs were rejected and zeroed out if 70% of the channels exceeded -3.5 to 5 standard deviations of a robust estimate of channel EEG power. This was completed using *clean_windows.m* (Kothe, 2010). Supplementary materials show that there was no violation of stationarity using this method and no bias towards any frequency.

EEG data were rejected based if more than 25% channels interpolated (42.5% of infant conditions and 25.5% of adult conditions removed); more than 25% continuous sections removed (14.9% vs 12.8% conditions removed for infant vs adult participants).

### 2.6. Data Analysis Plan

The first analysis investigated whether different low amplitude modulation frequencies cause different strengths of neural entrainment in infant and adult participants. This was completed by comparing the amplitude of the signal to noise ratio value at the target frequency bin across conditions using an ANOVAN. The subsequent analyses will only use the low amplitude modulated frequency which shows the strongest entrainment response.

The second analysis tested the sensitivity of power, phase and variance analyses using fast Fourier transform (FFT), phase locking value (PLV) and entropy of phase locking value (entropy) respectively. These will be completed on the participant samples and on individual participants.

The third analysis examined whether there is an increase or decrease in entrainment response over time. The four-minute condition was segmented into 2-minute, 1 minute and 30 second epochs. Power and phase analyses were completed to show trends of entrainment over time.

Finally, the fourth analysis studied entrainment responses using predetermined regions of interest from previous literature vs whole head neural responses.

#### 2.6.1. Analysis 1 – Between Low Amplitude Modulated Frequencies

Our first research question was to investigate whether different low amplitude modulation frequencies cause different strengths of neural entrainment in infant and adult participants. To address this, first for each participant x electrode x condition timeseries, a check was run to ensure that each timeseries was the same length. Any timeseries that was shorter than the required 122880 samples (512Hz x 240 seconds) were zero padded. A check was conducted to determine whether the zero padding would have an impact on the spectral composition, see supplementary materials. An FFT was calculated using a modified *myFFT.m* (Schoof, 2017) MATLAB script.

PLV (Lachaux et al., 1999) was calculated for each of the participant x electrode timeseries against frequencies of interest 2Hz above and below the condition’s target frequency in 0.01Hz steps. For example, for the 8Hz condition, a 6-10Hz window was selected giving 401 frequency bins. For the 2Hz condition, frequencies of interest were 1-4Hz in 0.01Hz steps due to the high pass filter completed during preprocessing stage. Any timeseries that was shorter than the required 122880 samples were zero padded in the same way as the FFT analysis. For each participant x electrode timeseries, a perfect sinusoid signal for each frequency bin within the frequencies of interest range was created with the same temporal length. A phase angle timeseries was calculated for the EEG timeseries and each of the frequencies of interest by taking the Hilbert transform of the data and calculating the phase angle. The difference between the neural phase angle timeseries and the frequencies of interest timeseries were taken, multiplied by the imaginary operator i to give a complex number and the exponential was determined for each value. PLV was calculated as the average exponential value in the timeseries. One PLV value is calculated for each frequency of interest.

After power (FFT) and phase (PLV) analyses were completed on the cleaned data sets in MATLAB 2021b (see Supplementary Materials for scripts used) the resulting data were then passed through a signal to noise ratio (SNR) script to standardise the data across conditions by removing the 1/f component of the neural data for each electrode x participant (e.g. Vettori et al., 2019).

SNR scores were calculated using a moving window of 25 frequency bins for each dataset between the first and last frequency of interest, for each participant x condition spectral series, in 0.01Hz steps. Of the 24 bins either side of the central frequency, the two closest frequency bins, and two bins within the remaining range with the maximum absolute difference from the target were removed and the average of the remaining 20 bins was taken from the central frequency bin.

For each condition x electrode x participant, the amplitude of the SNR score at the stimulation frequency was taken. An N-ways ANOVA (ANOVAN), *anovan.m,* was completed to test the significance of the SNR amplitude across stimulation frequency conditions (rest, 2Hz, 4Hz, 6Hz, 8Hz, 10Hz and 12Hz). An ANOVAN was chosen due to the uneven participant numbers per condition. Multiple comparisons, using *multcompare.m,* were completed when an ANOVAN showed significant differences.

#### 2.6.2. Analysis 2 – Sensitivity of power, phase and variance analyses

Our second research question tested the sensitivity of power, phase and variance analyses using FFT, PLV and entropy of phase locking value respectively. To investigate this, using only the stimulation frequency found to show strongest entrainment in analysis 1, FFT and PLV were calculated for each participant x electrode timeseries in the same way as in analysis 1.

Additionally, normalised entropy (Shannon, 1948; Tass et al., 1998) of PLV was calculated for each participant x electrode timeseries. Each timeseries was compared against frequencies of interest 1-4Hz in 0.01Hz steps. A phase angle timeseries for neural data and each sinusoidal signal for the frequencies of interest, along with the phase angle difference between the EEG timeseries and each of the frequencies of interest were created in the same way as in the PLV analysis. The entropy value was calculated over 80 bins (Notbohm, Kurths and Herrmann, 2016) using the below formula (Gross et al, 2021; Notbohm, Kurths, Herrmann, 2016), where H is the entropy and Pi is the probability of a given phase.

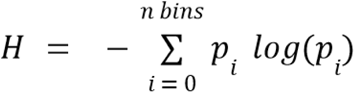

The probability of a given phase was calculated using *hist.m* to calculate the count and centre of each bin. The bin width was then determined as the difference between bin centres with *diff.m* and the maximum minus the minimum value. The counts per bin above 0 were collated and the probability of a value in each bin was calculated as the number of above zero values in the bin divided by the total number of values.

The resulting data were then passed through the same SNR script as in analysis 1. For the group wide analyses the data was averaged over electrodes and participants to give one FFT, PLV and entropy dataset. For individual participant analyses, the data were averaged over the electrodes only to give one data set for each of the analysis types per participant.

Next a permutation analysis was completed on each dataset. The SNR value at 2Hz was recorded as the test value. Bootstrapping with replacement was completed to create 10,000 surrogate datasets per participant and analysis type using *bootstrp.m* (Efron & Tibshirani, 1993). For each surrogate dataset the original test value was ranked against all surrogate values to give a test value rank. The 10,000 test value ranks generated with the surrogate datasets were then averaged to give an overall rank. The overall rank was considered significant if it fell in the top 5% of ranks.

#### 2.6.3. Analysis 3 – Entrainment over time analyses

Our third analysis examined whether there is an increase or decrease in entrainment response over time. To test this, using only the stimulation condition with the strongest entrainment, to see if there was an increase or decrease in entrainment over time, preprocessed data were segmented into 1 x 4 minute window, 2 x 2 minute windows, 4 x 1 minute windows or 8 x 30 second windows.

FFT and PLV were calculated for each participant x electrode x time window timeseries in the same way as in analysis 1. Subsequently SNR values were calculated in the same way as analysis 1.

SNR spectral series were averaged over participant and electrode to give one dataset for infants and adults, for both power and phase analysis, and for each time window. Then, permutation analysis was conducted in the same way as in analysis 2. This generated ranked permutation results in power and phase analyses for each time window and each participant group.

#### 2.6.4. Analysis 4 – Entrainment over space analyses

Our final analysis studied entrainment responses using five predetermined regions of interest from previous literature vs whole head neural responses. These five regions of interest were selected prior to the results being known. Electrodes referring to Cz; Fz-FCz-Fz; the central zenith line from anterior to posterior of the head (FPz, AFz, Fz, FCz, Cz, CPz, Pz, POz, Oz), the zenith line plus two lines of electrodes either side (referred to as expanded zenith line; Fp1, FPz, FP2, AF3, AFz, AF4, F1, Fz, F2, FC1, FCz, FC2, C1, Cz, C2, CP1, CPz, CP2, P1, Pz, P2, PO3, POz, PO4, O1, Oz, O2), and a square of 9 electrodes surrounding and including Cz (referred to as the vertex area; FC1, FCz, FC2, C1, Cz, C2, Cp1, CPz, CP2) were chosen as the potential regions of interest.

For each of these regions of interest, FFT and PLV were calculated for each participant x electrode for the full 4-minute timeseries in the same way as in analysis 1. Subsequently SNR values were calculated in the same way as analysis 1. SNR spectral series were averaged over participant and electrode cluster to give one dataset for infants and adults, for both power and phase analysis, for each region of interest. Then permutation analysis was conducted in the same way as in analysis 2. This generated ranked permutation results in power and phase analyses for each region of interest and each participant group.

Along with these analyses, topographic reporting of the whole head of electrodes was also completed and is shown in the results section to compare the ground truth regions of activation vs the clusters analysed.

## 3. Results

### 3.1. Analysis 1 – Between Stimulation Frequencies

A comparison of the entrainment to the low amplitude modulated frequency conditions rest (no sound), 2, 4, 6, 8, 10 and 12Hz was completed in both the power and phase domains using FFT and PLV.

#### 3.1.1. Power analysis

An ANOVAN comparing different SNR values of FFT results for rest (no sound) and stimulation frequencies (2, 4, 6, 8, 10 and 12Hz) for both infant (figure 3a) and adult (figure 3b) participants showed there was a significant difference between conditions in the power domain both for infants (F(6,63) = [40.996], p < 0.001) and adults (F(6,63) = [138.617], p < 0.001). There were no significant differences between electrodes (p > 0.05).

**Figure 1.**
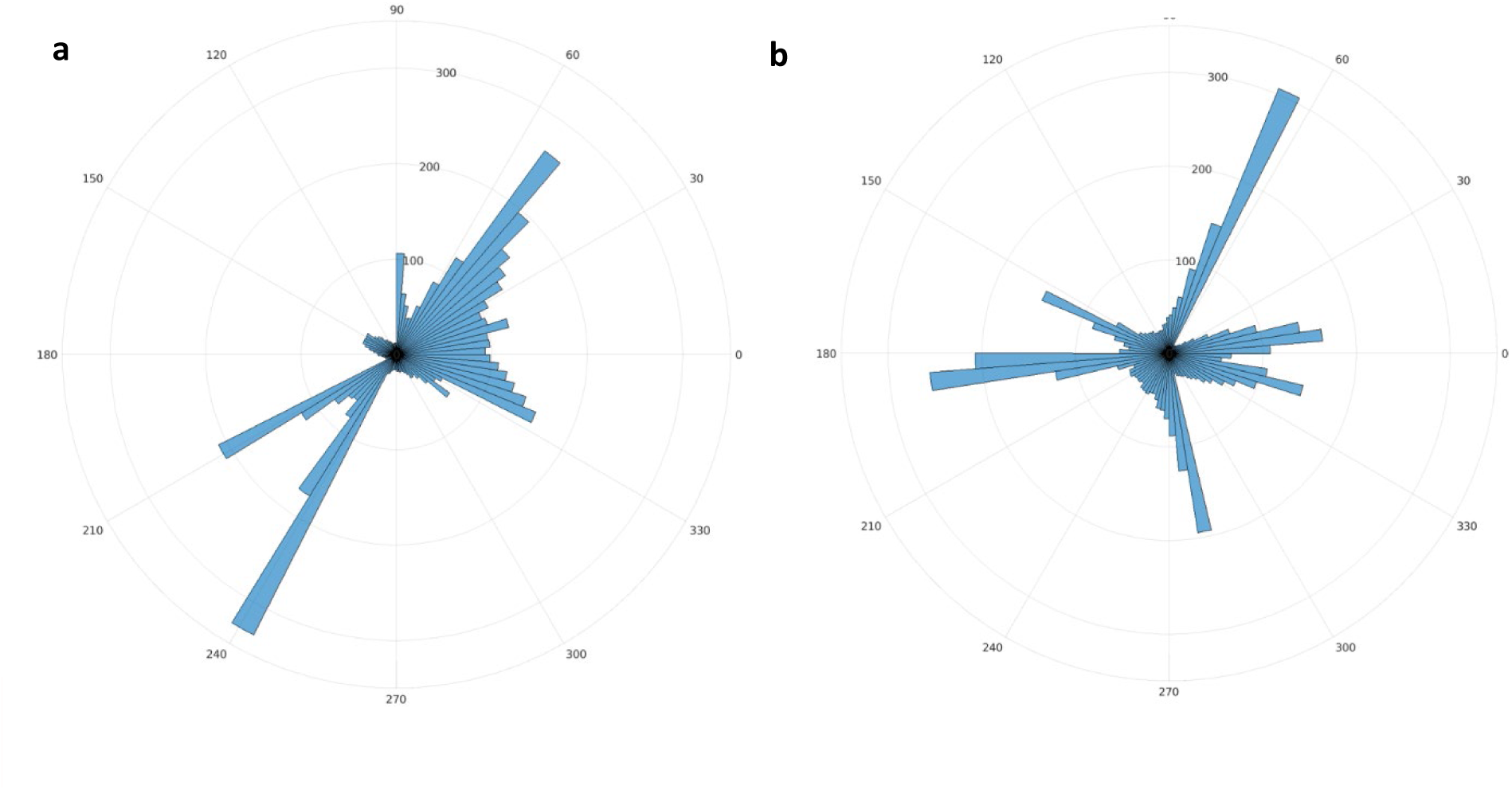
Examples of infant data that shows bipolar (a) and tripolar (b) phase responses to 8Hz stimuli. The large poles work to reduce the PLV value to a small response despite there beingphase angles shown on the polar plots that are clearly preferred.

**Figure 2.**
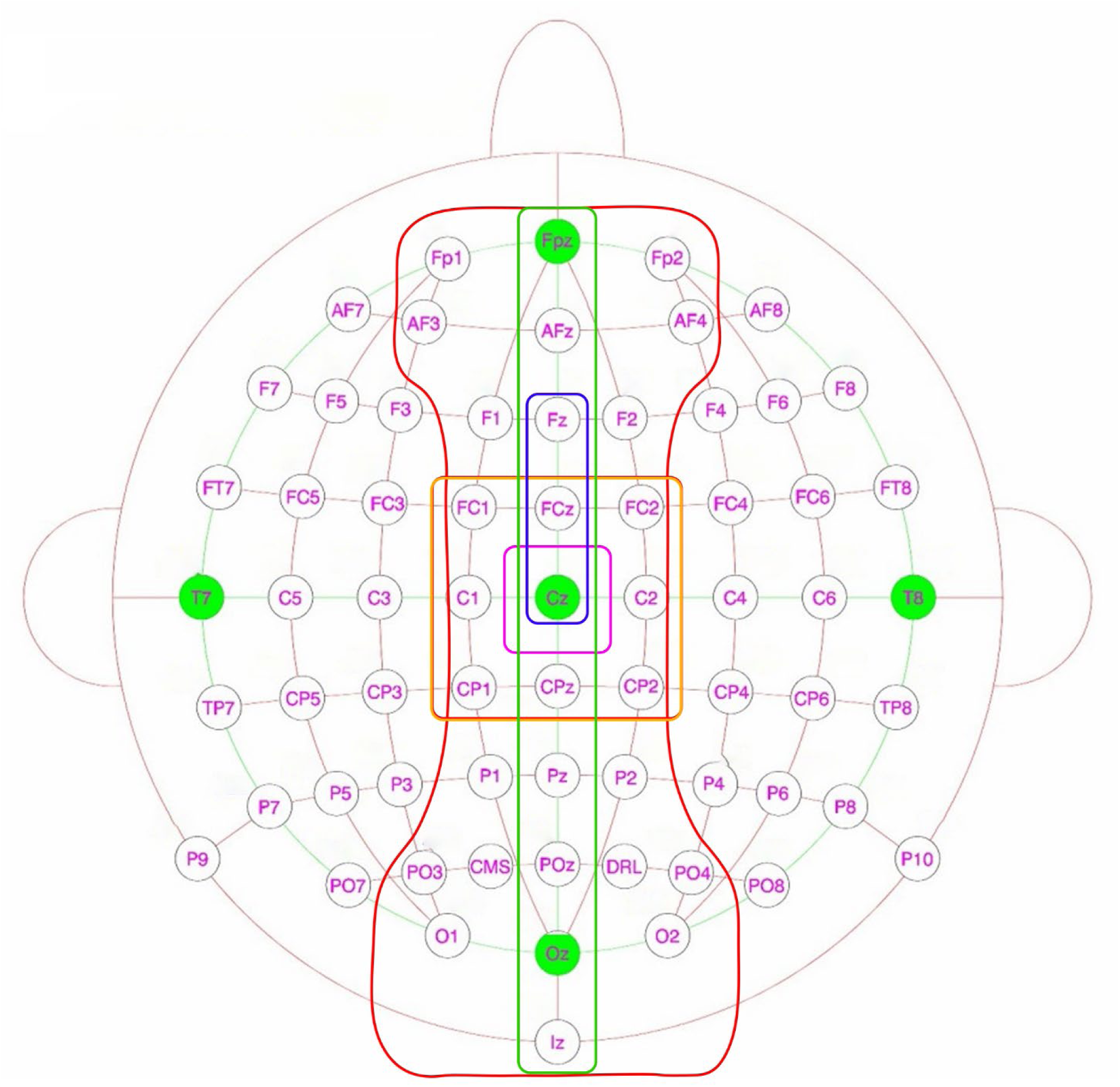
64 channel EEG cap using 10-20 montage, with highlighted areas relating to each region of interest chosen. Purple, Cz only; blue, Fz-FCz-Cz; orange, vertex area; green, zenith line; red, expanded zenith line.

**Figure 3.**
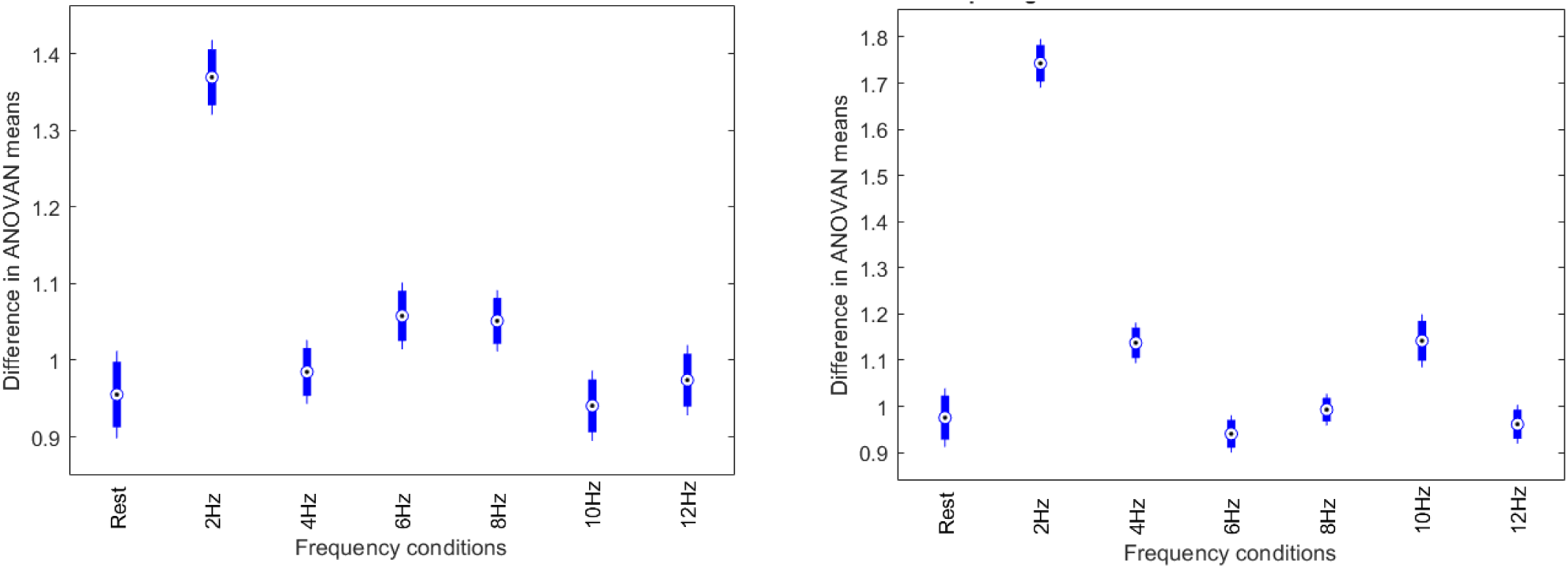
Multiple comparisons using multcompare.m between rest, 2, 4, 6, 8, 10 and 12Hz target frequency conditions, completed with an ANOVAN for infant (left) and adult (right) participants comparing SNR of FFT results.

Multiple comparisons were completed using the *multcompare.m* Matlab library, which showed that the 2Hz stimuli had the strongest response, being significantly higher than all other conditions for both infants and adults. Three other statistical differences were shown for infant participants (see table 1) between 10Hz and 6/8Hz and between 6Hz and rest. For adult participants, as well as differences between 2Hz and other frequency conditions, 4 and 10Hz conditions were also significantly different than rest, 6, 8 and 12Hz conditions. Statistical differences for adult participants are shown in table 1.

**Table 1.**
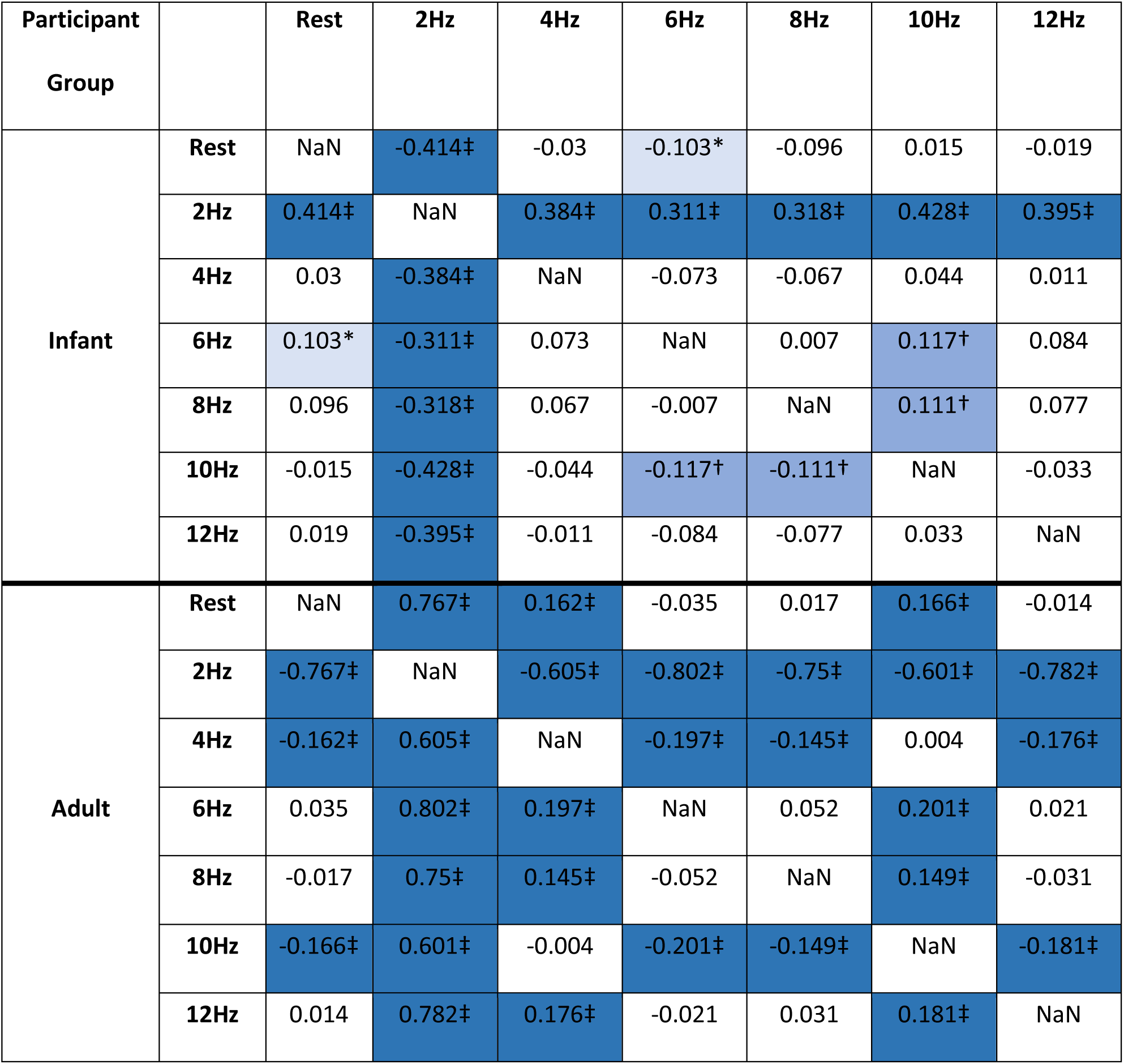
Differences of means between conditions for infant and adult SNR of FFT, dark blue and ‡ denotes p < 0.001, medium blue and † denotes p < 0.01 and light blue and * denotes p < 0.05.

| Participant Group |  | Rest | 2Hz | 4Hz | 6Hz | 8Hz | 10Hz | 12Hz |
| --- | --- | --- | --- | --- | --- | --- | --- | --- |
| Infant | Rest | NaN | -0.414‡ | -0.03 | -0.103* | -0.096 | 0.015 | -0.019 |
|  | 2Hz | 0.414‡ | NaN | 0.384‡ | 0.311‡ | 0.318‡ | 0.428‡ | 0.395‡ |
|  | 4Hz | 0.03 | -0.384‡ | NaN | -0.073 | -0.067 | 0.044 | 0.011 |
|  | 6Hz | 0.103* | -0.311‡ | 0.073 | NaN | 0.007 | 0.117† | 0.084 |
|  | 8Hz | 0.096 | -0.318‡ | 0.067 | -0.007 | NaN | 0.111† | 0.077 |
|  | 10Hz | -0.015 | -0.428‡ | -0.044 | -0.117† | -0.111† | NaN | -0.033 |
|  | 12Hz | 0.019 | -0.395‡ | -0.011 | -0.084 | -0.077 | 0.033 | NaN |
| Adult | Rest | NaN | 0.767‡ | 0.162‡ | -0.035 | 0.017 | 0.166‡ | -0.014 |
|  | 2Hz | -0.767‡ | NaN | -0.605‡ | -0.802‡ | -0.75‡ | -0.601‡ | -0.782‡ |
|  | 4Hz | -0.162‡ | 0.605‡ | NaN | -0.197‡ | -0.145‡ | 0.004 | -0.176‡ |
|  | 6Hz | 0.035 | 0.802‡ | 0.197‡ | NaN | 0.052 | 0.201‡ | 0.021 |
|  | 8Hz | -0.017 | 0.75‡ | 0.145‡ | -0.052 | NaN | 0.149‡ | -0.031 |
|  | 10Hz | -0.166‡ | 0.601‡ | -0.004 | -0.201‡ | -0.149‡ | NaN | -0.181‡ |
|  | 12Hz | 0.014 | 0.782‡ | 0.176‡ | -0.021 | 0.031 | 0.181‡ | NaN |

#### 3.1.2. Phase analysis

An ANOVAN comparing different SNR values of PLV results between stimulation frequencies for both infant (figure 4a) and adult (figure 4b) participants both show significant differences (infant, F(6, 63) = [35.903], p < 0.001; adult, F(6,63) = [131.33], p < 0.001). There were no significant differences between electrodes (p > 0.05).

**Figure 4.**
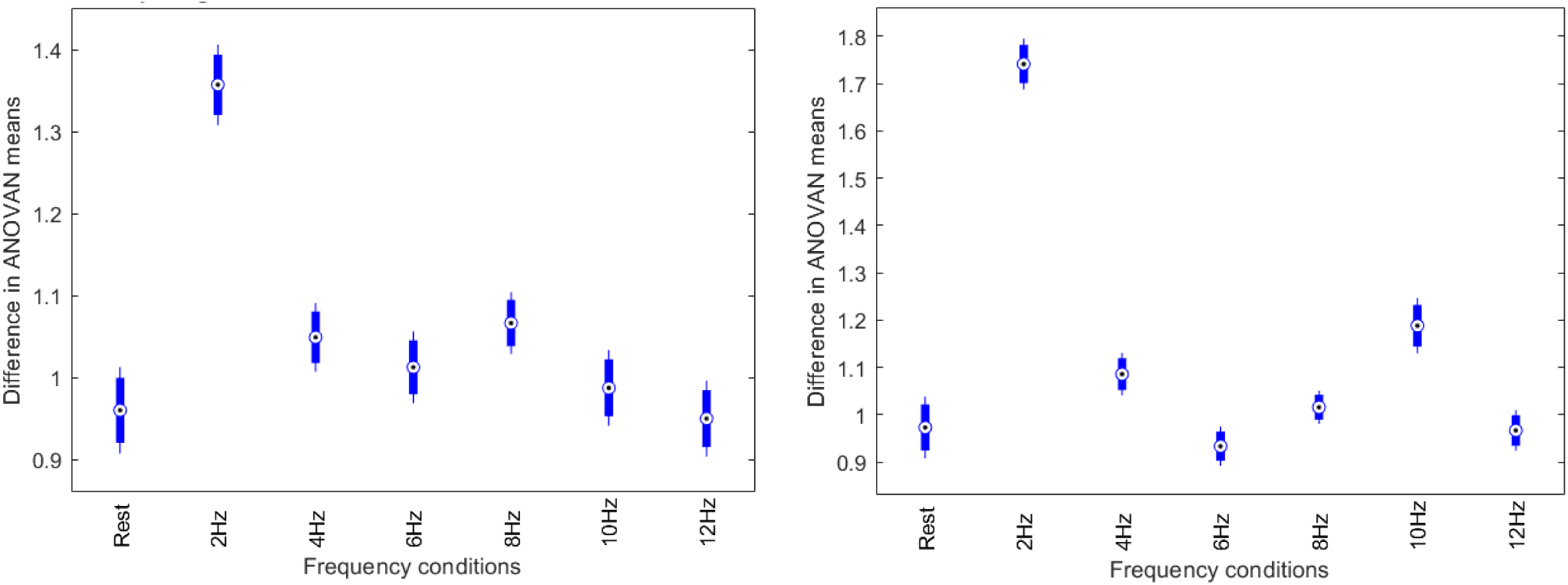
Multiple comparisons using multcompare.m between rest, 2, 4, 6, 8, 10 and 12Hz target frequency conditions, completed with an ANOVAN for infant (left) and adult (right) participants comparing SNR of PLV results.

Multiple comparisons were completed using the *multcompare.m* Matlab library, which showed that the 2Hz stimuli had the strongest response for both adults and infants, being significantly different from all other conditions. Three other statistically significant results were found, shown in table 2. For adult participants multiple other statistical differences were found, which are shown in table 2.

**Table 2.** Differences of means between the rest, 2, 4, 6, 8, 0 and 12Hz conditions for infant SNR of PLV dark blue and ‡ denotes p < 0.001, medium blue and † denotes p < 0.01 and light blue and * denotes p < 0.05.

| Participant Group |  | Rest | 2Hz | 4Hz | 6Hz | 8Hz | 10Hz | 12Hz |
| --- | --- | --- | --- | --- | --- | --- | --- | --- |
| Infant | Rest | NaN | 0.397‡ | 0.089 | 0.052 | 0.107† | 0.027 | -0.01 |
|  | 2Hz | -0.397‡ | NaN | -0.308‡ | -0.345‡ | -0.291‡ | -0.37‡ | -0.408‡ |
|  | 4Hz | -0.089 | 0.308‡ | NaN | -0.037 | 0.017 | -0.062 | -0.099* |
|  | 6Hz | -0.052 | 0.345‡ | 0.037 | NaN | 0.054 | -0.025 | -0.063 |
|  | 8Hz | -0.107† | 0.291‡ | -0.017 | -0.054 | NaN | -0.079 | -0.117† |
|  | 10Hz | -0.027 | 0.37‡ | 0.062 | 0.025 | 0.079 | NaN | -0.037 |
|  | 12Hz | 0.01 | 0.408‡ | 0.099* | 0.063 | 0.117‡ | 0.037 | NaN |
| Adult | Rest | NaN | 0.768‡ | 0.113* | -0.04 | 0.043 | 0.215‡ | -0.006 |
|  | 2Hz | -0.768‡ | NaN | -0.655‡ | -0.808‡ | -0.725‡ | -0.553‡ | -0.774‡ |
|  | 4Hz | -0.113* | 0.655‡ | NaN | -0.152‡ | -0.07 | 0.102 | -0.119† |
|  | 6Hz | 0.04 | 0.808‡ | 0.152‡ | NaN | 0.082* | 0.255‡ | 0.033 |
|  | 8Hz | -0.043 | 0.725‡ | 0.07 | -0.082* | NaN | 0.172‡ | -0.049 |
|  | 10Hz | -0.215 | 0.553‡ | -0.102 | -0.255‡ | -0.172‡ | NaN | -0.221‡ |
|  | 12Hz | 0.006 | 0.774‡ | 0.119† | -0.033 | 0.049 | 0.221‡ | NaN |

### 3.2. Analysis 2 – Sensitivity of power, phase and variance analyses

Having identified that the strongest entrainment between conditions was observed at 2Hz, we went on to investigate the sensitivity of power, phase and variance analyses using FFT, PLV and entropy at both the group and individual level.

#### 3.2.1. Comparison of FFT, PLV and entropy at a group level

A bootstrapping analysis of the SNR scores derived from FFT, PLV and entropy analyses was conducted. Figure 5 shows the spectral series for adults and infants for each analysis type. Table 3 shows the SNR values for each participant and analysis condition. All conditions were highly significant (p < 0.01) except for infant entropy (p > 0.05).

**Figure 5.**
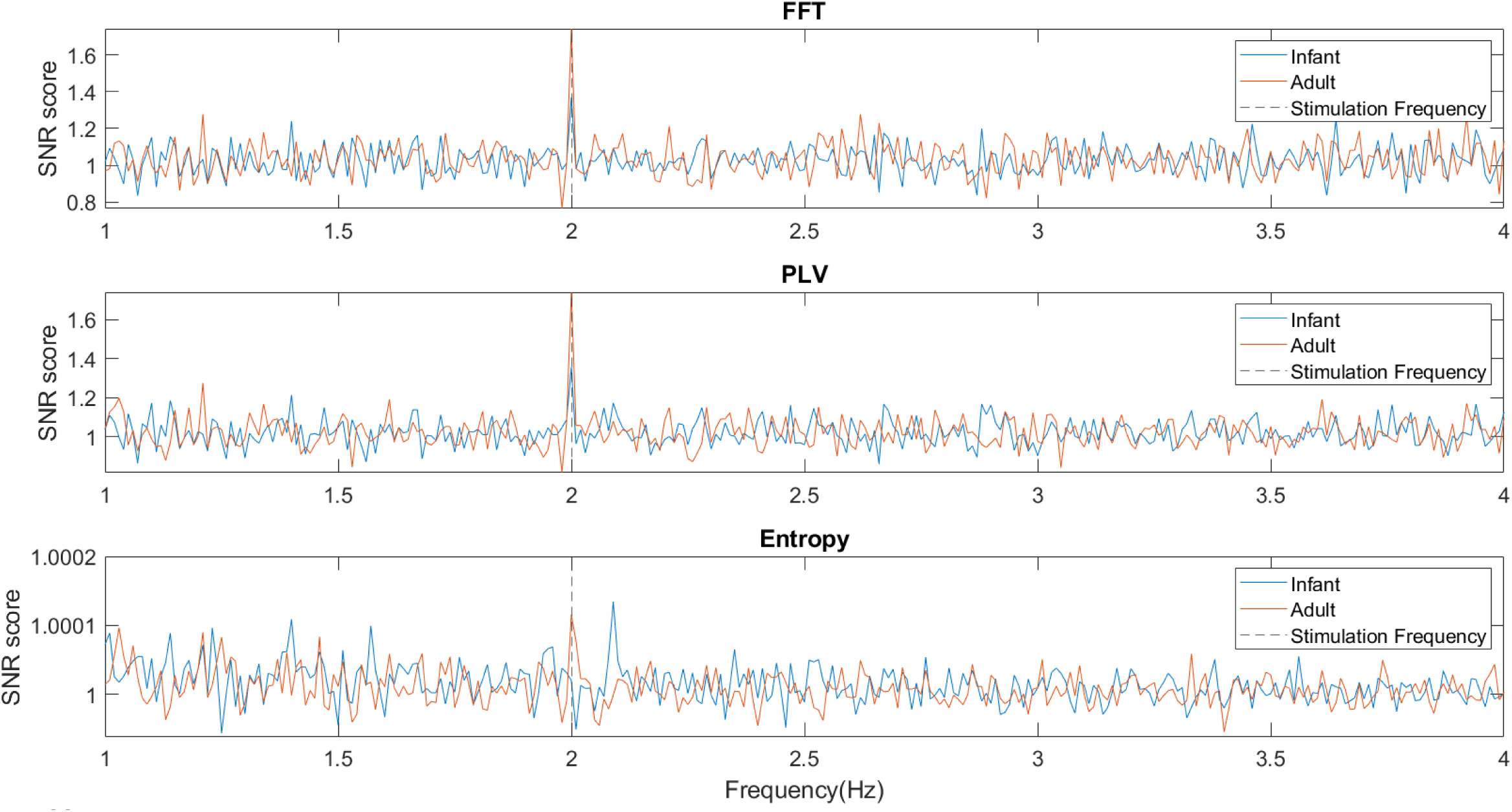
SNR scores for each of the analysis types, from 1-4Hz with a frequency resolution of 0.01Hz and a 2Hz stimulation frequency. Adults showed a strong response compared to other frequencies in all analysis types. Infants showed a response for FFT and PLV analyses that was weaker than adults and showed no response at the stimulation frequency for the entropy of PLV analysis.

**Table 3.**
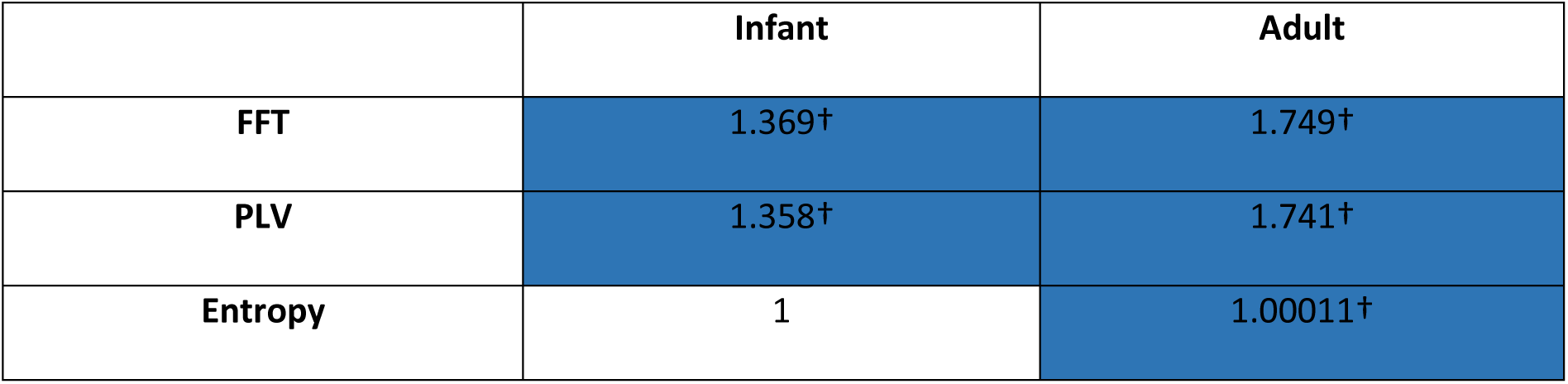
Mean SNR values at the stimulation frequency of 2Hz for infant and adult participant groups for each analysis type (FFT, PLV, entropy of PLV). dark blue and † denotes p < 0.01.

|  | Infant | Adult |
| --- | --- | --- |
| **FFT** | 1.369† | 1.749† |
| **PLV** | 1.358† | 1.741† |
| **Entropy** | 1 | 1.00011† |

To investigate if there was a difference between infant and adult participant populations, a Welch’s t-test was completed between infants and adult SNR scores at the 2Hz stimulation frequency for each analysis type to test for differences between the participant groups. This was not significantly different for any of the groups (p > 0.05). An ANOVA comparing the mean ranks achieved for each participant’s bootstrapped values between each analysis type was completed to investigate if the results were significantly different between analysis types. This was not significant for either infants or adults (p > 0.05).

#### 3.2.2. Comparison of FFT, PLV and entropy at an individual level

To investigate individual differences in frequency tagging responses and examine if the significant findings at the group level were driven by a small number of individuals or were seen across the group, analyses on a participant-by-participant basis were conducted. The analysis was the same as at the group level (section 3.2.1) but used datasets that had not been averaged over participants. Table 4 shows the per participant data and significance. Data quality metrics per participant are shown in Supplementary Materials.

**Table 4.** Mean SNR values per participant (both infant or adult) for each analysis type (FFT, PLV, entropy of PLV), black shading indicates datasets that were not present due to technical error or poor data quality, medium blue and † denotes p < 0.01 and light blue and * denotes p < 0.05

|  | Infant |  |  | Adult |  |  |
| --- | --- | --- | --- | --- | --- | --- |
|  | FFT | PLV | Entropy | FFT | PLV | Entropy |
| 1042 | 1.196 | 1.286 | 1 | 2.267* | 2.299† | 1.0001† |
| 1051 | 1.529 | 1.834† | 0.99994 | 1.305* | 1.122 | 1.0001 |
| 1052 | 1.323 | 1.508* | 1.0001 |  |  |  |
| 1053 | 0.932 | 0.788 | 0.99989 | 1.671* | 1.592† | 1.0001* |
| 1055 |  |  |  | 1.847† | 1.806† | 1.0001 |
| 1059 |  |  |  | 1.301 | 1.22 | 1 |
| 1061 |  |  |  | 1.804† | 1.648† | 1.0002† |
| 1062 | 1.271 | 0.999 | 0.99993 |  |  |  |
| 1068 |  |  |  | 2.006† | 2.503† | 1.0002† |
| 1079 | 2.058† | 1.89† | 1.0002* |  |  |  |
| 1082 | 1.096 | 1.011 | 0.99993 |  |  |  |
| 1084 | 1.548* | 1.545† | 1.0001 |  |  |  |

There were more adult participants that were found to have a significant frequency tagging response at 2Hz (FFT, 86%; PLV 71%; entropy 57%) compared to infant participants (FFT, 25%; PLV 50%, entropy 13%). There were also more highly significant results for adult participants across analyses (52%) compared to infant results (17%).

### 3.3. Analysis 3 – Entrainment over time analyses

The 2Hz stimulation condition was the only condition used, as this had shown the strongest entrainment response (see analysis 1). To investigate if there is a progression or reduction in entrainment as measured by different analyses, a comparison of four-time lengths was made for power (FFT) and phase (PLV) analyses. Conditions contained 4-minute recordings which were split into either: 1 x 4-minute windows, 2 x 2-minute windows, 4 x 1-minute windows or 8 x 30-second windows. A bootstrapping analysis was used to compare the test value derived from the 2Hz frequency bin for each of the temporal epochs in the same way as in analysis 2. Table 5 shows the SNR score at the 2Hz stimulation frequency across segments for each participant group, analysis type and time segments.

**Table 5.** Averaged SNR value at the 2Hz stimulation frequency split by participant group, analysis and time window. † denotes p < 0.01 and * denotes p < 0.05.

| Participant group | Analysis | Time window | Test values |  |  |  |  |  |  |  |
| --- | --- | --- | --- | --- | --- | --- | --- | --- | --- | --- |
| Infant | FFT | 4 minutes | 1.369† |  |  |  |  |  |  |  |
|  |  | 2 minutes | 1.213* |  |  |  | 1.265† |  |  |  |
|  |  | 1 minute | 1.214* |  | 1.180* |  | 1.184* |  | 1.034 |  |
|  |  | 30 seconds | 1.164* | 0.988 | 1.052 | 1.194* | 1.050 | 1.116 | 1.089 | 1.016 |
|  | PLV | 4 minutes | 1.358† |  |  |  |  |  |  |  |
|  |  | 2 minutes | 1.251* |  |  |  | 1.197† |  |  |  |
|  |  | 1 minute | 1.218† |  | 1.168* |  | 1.156* |  | 1.033* |  |
|  |  | 30 seconds | 1.161* | 0.998 | 1.048 | 1.138 | 1.059 | 1.143* | 1.055 | 0.993 |
| Adult | FFT | 4 minutes | 1.743† |  |  |  |  |  |  |  |
|  |  | 2 minutes | 1.358† |  |  |  | 1.421† |  |  |  |
|  |  | 1 minute | 1.208* |  | 1.276† |  | 1.388† |  | 1.216* |  |
|  |  | 30 seconds | 1.038 | 1.072 | 0.994 | 1.243 | 1.410† | 1.168* | 1.194* | 1.108 |
|  | PLV | 4 minutes | 1.742† |  |  |  |  |  |  |  |
|  |  | 2 minutes | 1.381† |  |  |  | 1.469† |  |  |  |
|  |  | 1 minute | 1.219† |  | 1.266† |  | 1.389† |  | 1.195* |  |
|  |  | 30 seconds | 1.020 | 1.075 | 1.006 | 1.198† | 1.211† | 1.205* | 1.147* | 1.112 |

### 3.4. Analysis 4 – Entrainment over space analyses

To test the spatial location of the frequency tagging response in both infants and adults, and to test the validity of regions of interest selected in previous research. A bootstrapping permutation analysis was conducted that was the same as in analysis 2 and applied to each region of interest (electrode locations are listed in section 2.6.4). Table 6 shows which areas of interest showed significance, with all areas being significant for both FFT and PLV for adults. In infant data Fz-FCz-Cz, zenith line and expanded zenith line were significant in the FFT data (p < 0.01, p < 0.05, p < 0.05) and Fz-FCz-Cz, expanded zenith line and vertex area were significant in the PLV data (all p < 0.05).

**Table 6.** Showing averaged SNR values at 2Hz for each region of interest per analysis and participant type. medium blue and † denotes p < 0.01 and light blue and * denotes p < 0.05.

|  | SNR of FFT |  | SNR of PLV |  |
| --- | --- | --- | --- | --- |
|  | Infant | Adult | Infant | Adult |
| **All electrodes** | 1.369† | 1.749† | 1.358† | 1.741† |
| **Cz only** | 1.285 | 1.671† | 1.251 | 1.781† |
| **Fz-FCz-Cz** | 1.374† | 2.113† | 1.365* | 2.115† |
| **Zenith line** | 1.246* | 1.815† | 1.174 | 1.721† |
| **Expanded zenith line** | 1.234* | 1.817† | 1.215* | 1.797† |
| **Vertex area** | 1.232 | 1.789† | 1.220* | 1.818† |

To compare these regions to the significant clusters seen across the whole head, topoplots of SNR scores for FFT and PLV analyses for participant groups are shown in figure 6 (a-d), and significance values are mapped with the topoplots in figure 6 (e-h).

**Figure 6.**
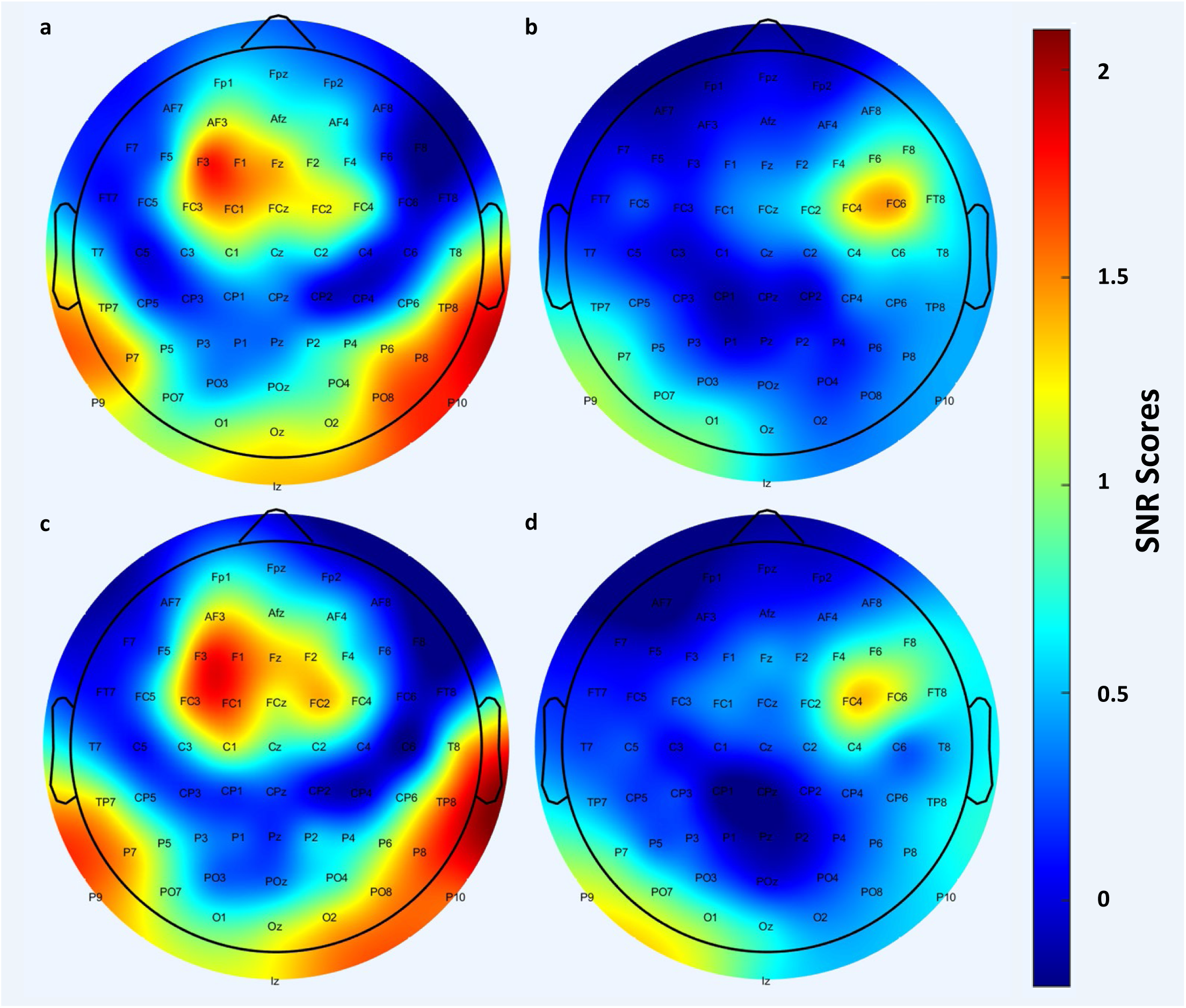
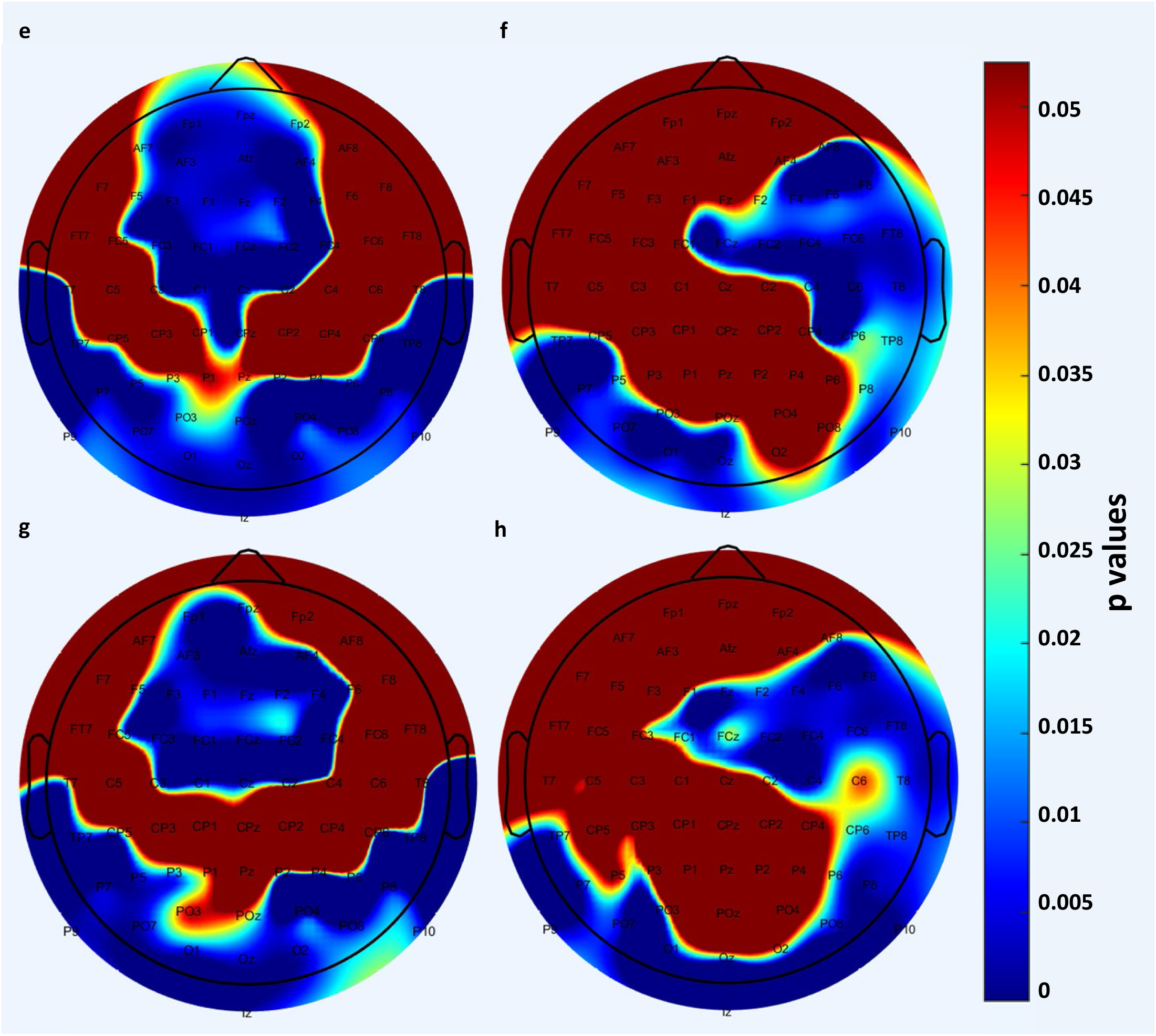
Topoplots showing the signal to noise ratio value at 2Hz (a-d) and areas of significance (e-h) p > 0.05 shown in red, p < 0.05 to p < 0.001 shown in various shades of blue as shown by the colour bar. Topoplots show a standard 64 channel 10-20 montage. Adult data shown on the left (a, c, e, g), infant data shown on the right (b, d, f, h), FFT data shown in plots a, b, e and f while PLV data is shown in plots c, d, g and h.

## 4. Discussion

### 4.1. Overview

In this study we examined how a range of low amplitude modulated frequencies impact the level of neural entrainment recorded with young (4–6-month-old) infants and their adult caregivers. We also investigated the sensitivity of analysis methods and spatiotemporal characteristics of infant and adult entrainment to the stimulation frequency that showed the strongest neural entrainment. To the best of our knowledge, no previous research has demonstrated a direct comparison of these important low amplitude modulated frequencies under the same conditions.

One issue when comparing across frequencies is the 1/f nature of neural data makes it difficult to compare absolute values in response to different stimulation frequencies (e.g. Cellier et al., 2021). Another issue is that the range of values expected for each analysis is different (e.g. Gross et al., 2021). To combat this, all results were subjected to a signal to noise ratio calculation to remove the average of the surrounding data and normalise the results. Below when referring to different frequency or analysis conditions these are all in fact the signal to noise ratio versions of these data so that they can be compared.

### 4.2. Between Low Amplitude Modulated Frequencies

The results comparing low amplitude modulated frequency conditions found that both adult and infant participants showed the strongest response in both the power and phase domains to the 2Hz stimuli when compared to all other conditions. While many stimulation frequencies have been used in a range of auditory frequency tagging studies (see section 1.4), to our knowledge this is the first study to directly compare this range of low amplitude modulated stimulation frequencies as a measure of infant and adult neural entrainment and shows that not all low amplitude modulated stimulation frequencies are responded to equally.

This contradicts our hypothesis that there would be an increased response to neural frequencies related to the resting states of the infant (6-9Hz) and adult (9-12Hz) brains. This was thought to be the case due to the increased power at these frequencies and previous research by Notbohm, Kurths and Sack, (2016) which demonstrated that increasing spectral distance between a neural oscillator and stimulus frequency requires an increase in the stimulus intensity to bring the two oscillators into alignment. While there was some increase in the neural entrainment response seen at these dominant frequencies to the stimulation frequency, this was dwarfed by the response to the 2Hz condition.

Aside from the 2Hz condition vs other stimulation frequencies there were some significant differences seen, which often did reflect the difference between frequencies that were at the participants’ dominant frequency vs others. In the power domain, for adult participants there were significant differences found with an increased response at 4 and 10Hz vs rest, 6, 8 and 12Hz frequencies. Whereas, in the infant group there were significant differences found with an increased response shown at 6 and 8Hz vs 10Hz, as well as 4Hz vs rest. In the phase domain, for adult participants, there was a significantly increased response from the 4 and 10Hz conditions when compared to rest, 6 and 12Hz conditions as well as an elevated response in 10Hz compared to 8Hz, and 8Hz compared to 6Hz conditions. For infant participants, there were significantly increased neural responses to the 4 and 8Hz conditions vs 12 Hz, as well as 8Hz vs rest.

This reinforces previous work (e.g., Kabdebon et al., 2022) that has suggested it is best to avoid stimulation frequencies that correspond to these dominant neural frequencies without an appropriate control. This could be an issue when comparing between groups of different ages that are above and below the ages of ∼7 years old. As noted in the paper by Cellier et al., (2021) the resting dominant frequencies of children changes at ∼7 years old from a dominant theta band of ∼7Hz to a dominant alpha band of ∼10Hz. These findings also present an issue for studies that choose to present frequency tagged stimuli at multiple frequencies which include dominant neural frequencies, without having an appropriate control. Researchers may not be able to differentiate how much of the result in this range is due to the frequency tagging condition and how much is due to an underlying endogenous oscillatory activity.

It is interesting that both infants and adults responded more strongly to the lowest amplitude modulated stimulation that was presented, which raises further questions. While 2Hz was the stimulation frequency that showed the strongest response in this study, it is not clear from these results whether this is because 2Hz condition truly shows the strongest response of all possible low amplitude modulated frequencies, or whether this shows the strongest response because this study didn’t investigate the 0.1-4Hz frequency range in enough detail. Future studies should investigate the neural response to stimuli between 0.1 and 4Hz to further our understanding.

The researchers of the current study considered confounding explanations for the strength of neural response at 2Hz. Perhaps the result at 2Hz represents the resting heart rhythm of the infants and adults, who have an average heart rhythm of 2.15Hz and 1.2Hz respectively (Ostchega et al., 2011). Similarly, it could be questioned whether the results are due to the 1/f distribution of the neural signal. However, for both of these theories, this would also have been seen in the resting data collected.

Another possibility is that the result could be attributed to the language heard by participants, English has many elements that follow a 2Hz structure including a 2Hz “stress” rate (Goswami and Leong, 2013; Leong, 2014), and infant directed speech also shows similar prosodic stress components at 2Hz (Leong and Goswami, 2015). While not all participants were native English speakers, all certainly had proficient English enough to live in the UK and it is to be expected that the infants will have been heavily exposed to English in their environments. Further studies are required in cultures that do not share this property with the English language.

### 4.3. Sensitivity of power, phase and variance analyses

On a group level, investigation of FFT, PLV and entropy analyses as a method of detecting frequency tagging responses showed that FFT and PLV demonstrated a significant frequency tagged neural response at the target stimulation frequency of 2Hz. This is consistent with previous studies (see section 1.5.3). Entropy showed the same result for adult participants only, suggesting that the variance in PLV as measured in the entropy measure was not a strong predictor of frequency tagged results. This contrasts with previous studies looking at entropy of PLV as a method to detect neural frequency tagged response (Notbohm, Kurths and Sack, 2018). This may be because previous studies have focused on entropy as a measure of frequency tagging in adults rather than young infants.

There were no significant differences found between infant and adult results at the 2Hz stimulation frequency for FFT or PLV analyses, but there was a significant difference found between infant and adult entropy values, which was to be expected.

On a participant-by-participant level, the results showed that not all participants had a significant frequency tagging response at 2Hz and that a significant result using one analysis did not always translate to a significant result across all analysis types. Adult participants had more individual participants that showed a frequency tagging response than infants. This likely caused the non-significant but lower response at 2Hz from infants as seen in figure 5. Individual participant details relating to age, infant gestation and gender are shown in supplementary materials. Infant participants that showed a neural response were not of one age group, gender or having had a particularly long or short gestation period. Data quality metrics are also shown in the supplementary materials. Analyses suggested that data quality was not a significant driver of whether significant entrainment was observed.

### 4.4. Entrainment over time analyses

To investigate whether there is an increase or decrease in entrainment as measured by the frequency tagging response, data were segmented into 4, 2 or 1 minute or 30 second chunks and bootstrapping analyses were conducted on each segment for each participant group. For infant FFT analysis, data remained significant throughout the 4- and 2-minute segments and for the first three of the 1-minute segments. Data were also significant for the 1^st^ and 4^th^ of the 30 second chunks. For infant PLV analysis, data were significant for all the 4-, 2- and 1-minute segments and the 1^st^ and 6^th^ 30 second segments. Infants showed no clear pattern of significant segments. Adult results showed all the 4-, 2- and 1-minute segments for both FFT and PLV analyses were significant as well as the 5- 7^th^ and 4-7^th^ segments in FFT and PLV analyses respectively. There was no clear pattern showing a change of significance results between the timeseries.

As the data for these segments is the same but divided into smaller and smaller pieces, it could be suggested that taking the spectral series from a smaller timeseries decreases the response seen to the stimulation frequency, which in turn reduces the likelihood of seeing a statistically significant result. This is evidenced by smaller segments of highly significant epochs not being subsequently significant, for example when comparing the first two 1 minute segments of adult FFT results (p < 0.05 and p < 0.01) to the first four 30 second segments (all p > 0.05). This suggests that the length of the stimulation is important to seeing a statistically significant result.

### 4.5. Spatial analyses

Previous studies have used specific regions of interest to investigate frequency tagging. These areas are often chosen to satisfy hypothesis driven research, but in many cases are also centred around auditory regions of the brain. Five ROIs were found in previous literature (see section 1.6), these were each tested to show if they would demonstrate a frequency tagging response. The ground truth frequency tagging response was also shown in topoplots. In adult data all five of the regions of interest showed significant results, while infant data showed significant results in the: Fz- FCz-Cz electrodes (FFT and PLV); zenith line region (FFT only); expanded zenith line (FFT and PLV) and in the vertex region (PLV only).

Further data showed that there were significant clusters in adult data in a slightly left lateralised fronto-central region and in a tempero-occipital cluster spread across both hemispheres. In infant data a right lateralised fronto-central-parietal cluster was shown along with a left lateralised occipital-temporal cluster. This may suggest that while there is some overlap between the predefined clusters that previous studies have used, there are other areas that are of interest outside the usual regions of interest. The lateralisation seen in infant data and the slight lateralisation on the opposite side of the adult data is also interesting as it may represent a developmental change in response to these stimuli. While no developmental conclusions can be drawn from the current results it is worth further study to see if there is a gradual or sudden shift in significant regional clusters over time.

## Conclusion

Careful choice of low amplitude modulated stimulation frequency should be an important consideration when designing a frequency tagging experiment. Based on current findings choosing a 2Hz low amplitude modulated frequency offers a comparable stimulation frequency between young infant and adult participants. Other stimulus and analysis choices are also worth considering, including having sufficient stimulus length and conducting whole head recordings where possible.

There were no significant differences between power and phase analyses.

## Data and Code Availability

MATLAB code, stimuli files, raw EEG files and EEG events used are freely available [here](https://e1.pcloud.link/publink/show?code=kZlkQYZ2Jjab25UEGu8WcGTTXqvMXG5dVmV). Participant details are available upon request from the corresponding author due to the potentially sensitive nature.

## Ethics Statement

The University of East London ethics committee approved the study (application ID: ETH2021-0076). All adult participants provided informed consent for both themselves and their children according to the Declaration of Helsinki. All participants were offered a £10 shopping voucher as a monetary reward for their time. Travel and food expenses were also covered for those that requested them.

## Acknowledgements

This project was funded by the European Research Council (ERC Horizon 2020; award number 853251). The authors thank Dr Sofie Vettori and Dr Victoria Leong for their helpful comments during the initial stages of this article.

## Supplementary materials

### Participant details

Blanks indicate where the adult participant did not give an answer.

| Ppt No | Adult age (years) | Infant age (days) | Infant gender | Inf gestation (weeks) | Other children |
| --- | --- | --- | --- | --- | --- |
| 1009 | 38.36 | 155 | M | 44 | Y |
| 1028 | 32.84 | 153 | F | 38.5 | N |
| 1039 | 33.70 | 150 | M | 40 | N |
| 1040 | 35.59 | 144 | M |  | Y |
| 1046 | 40.15 | 184 | M | 42 | Y |
| 1051 | 32.81 | 159 | M | 44 | N |
| 1052 | 32.30 | 157 | F | 47 | N |
| 1053 | 34.86 | 155 | M | 39 | Y |
| 1055 | 34.52 | 161 | F | 38 | Y |
| 1059 |  | 178 | M |  |  |
| 1060 |  | 174 | F |  |  |
| 1061 | 42.42 | 178 | F | 38 | Y |
| 1062 |  | 172 | F |  |  |
| 1064 | 31.80 | 161 | M | 41 | N |
| 1066 | 34.55 | 163 | M | 39 | N |
| 1068 | 41.04 | 151 | M | 39 | Y |
| 1070 |  | 166 | F |  |  |
| 1076 | 35.95 | 190 | F |  |  |
| 1077 | 35.70 | 138 | M | 39 | N |
| 1078 | 34.33 | 181 | M | 38 | N |
| 1079 | 33.57 | 136 | F | 40 |  |
| 1082 | 29.79 | 194 | M |  |  |
| 1084 | 39.52 | 173 | M | 42.1 |  |
| 2069 | 39.21 | 174 | M | 40 |  |
| 3042 | 35.37 | 153 | M | 37 | N |
| 4032 | 33.20 | 177 | F | 44 | N |

Conditions completed (post preprocessing)

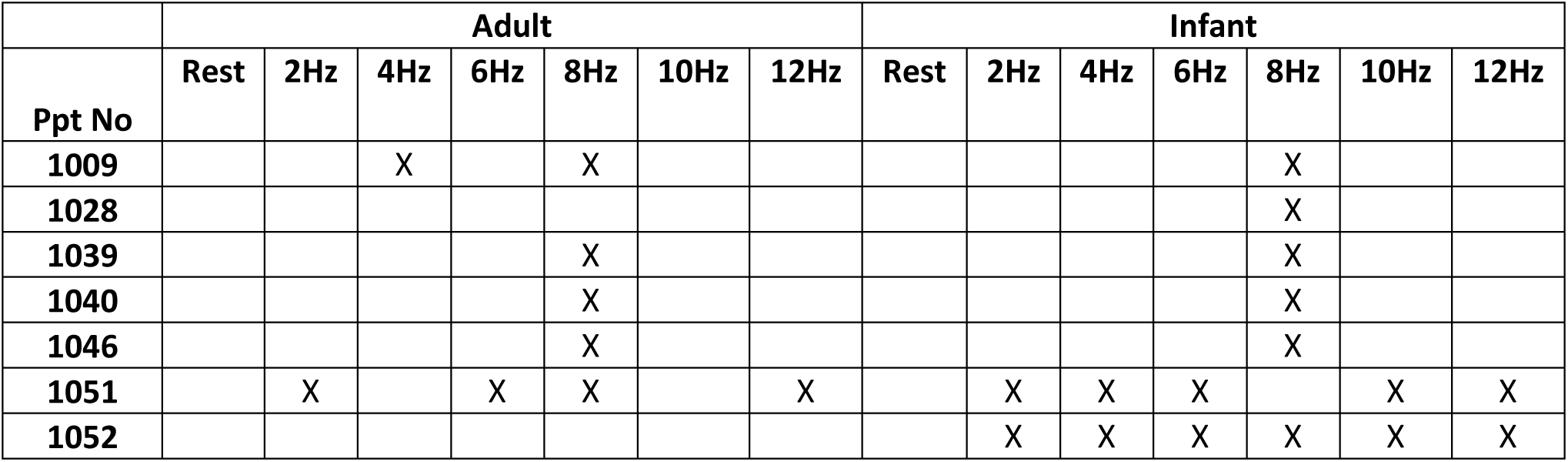

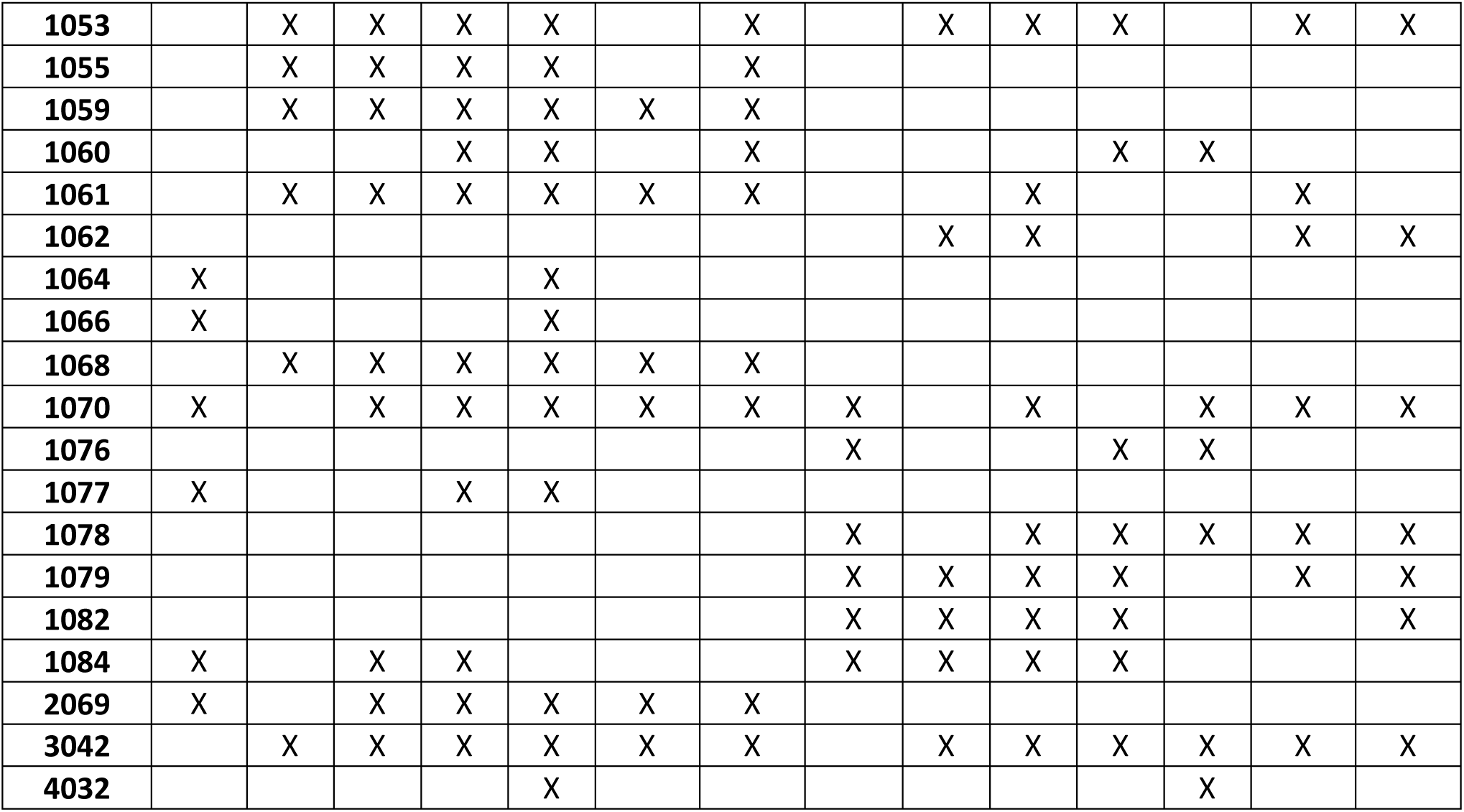

Check for impact of zero padding the results

In MATLAB, a composite sinewave was produced by adding a series of sinewaves with amplitude modulated frequencies of 1-100 + 10%, i.e. 1.1, 2.2, 3.3 through to 110, the time series can be seen in the below figure (a), along with a close up (b) and the spectral series (c). Then for each one second segment a random number of segments was zeroes out (up to 25% which is the maximum threshold in this study), which can be seen in the spectral (d) and temporal (e) series.

While there was a reduction in amplitude of the recorded frequencies between c and e there was no distortion of the spectral make up of the signal. There was some added low level noise broadband across the spectra.

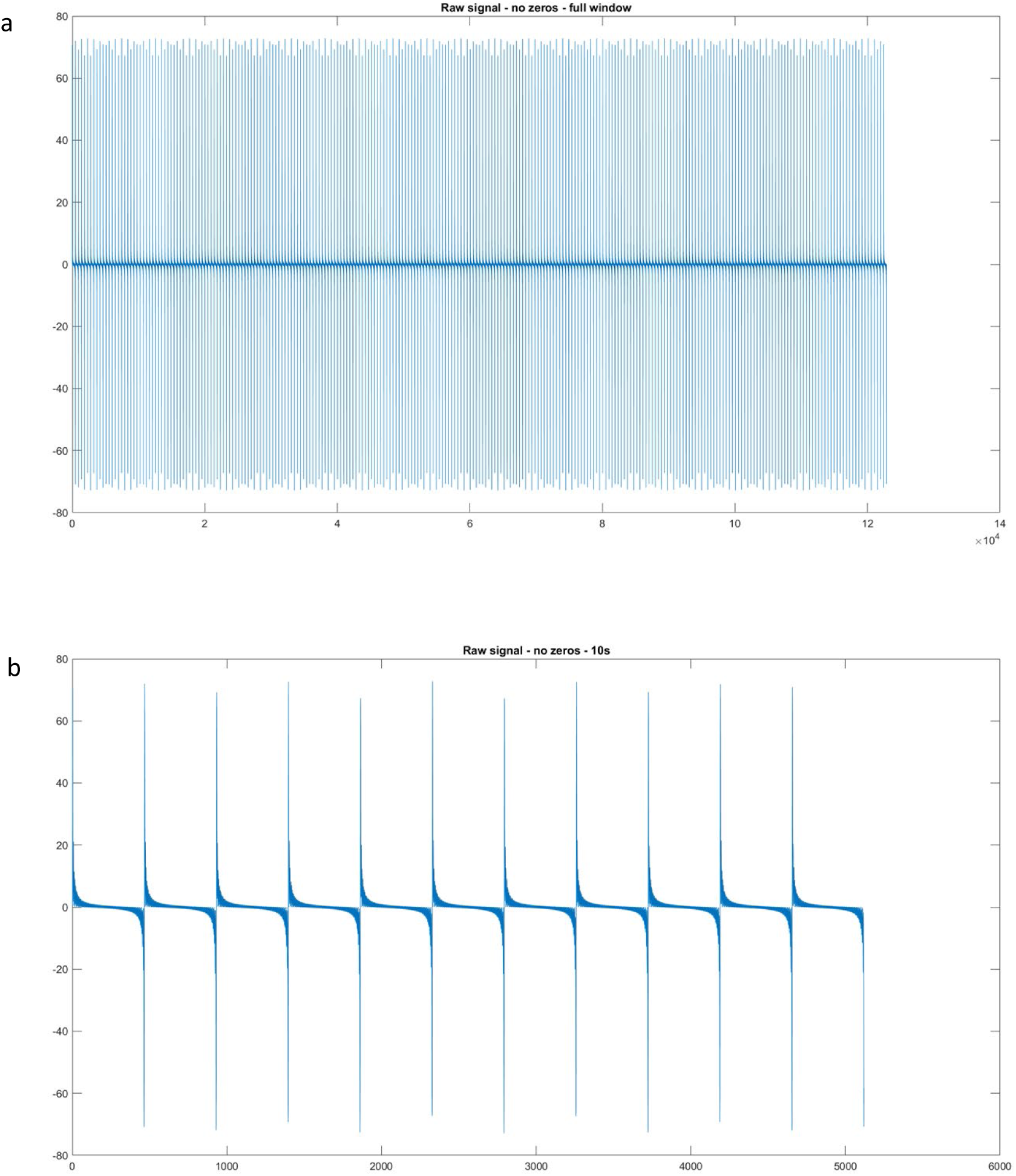
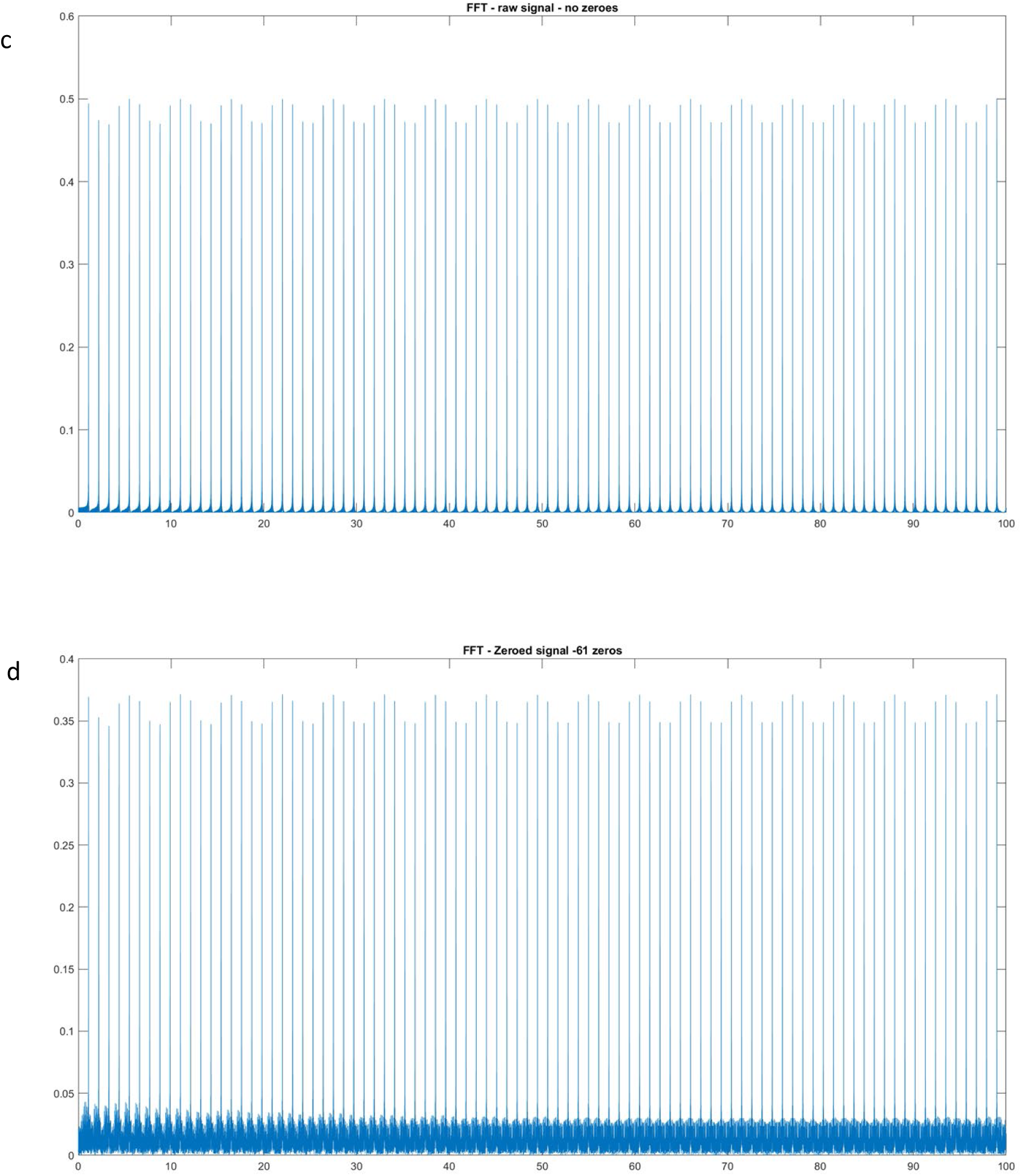
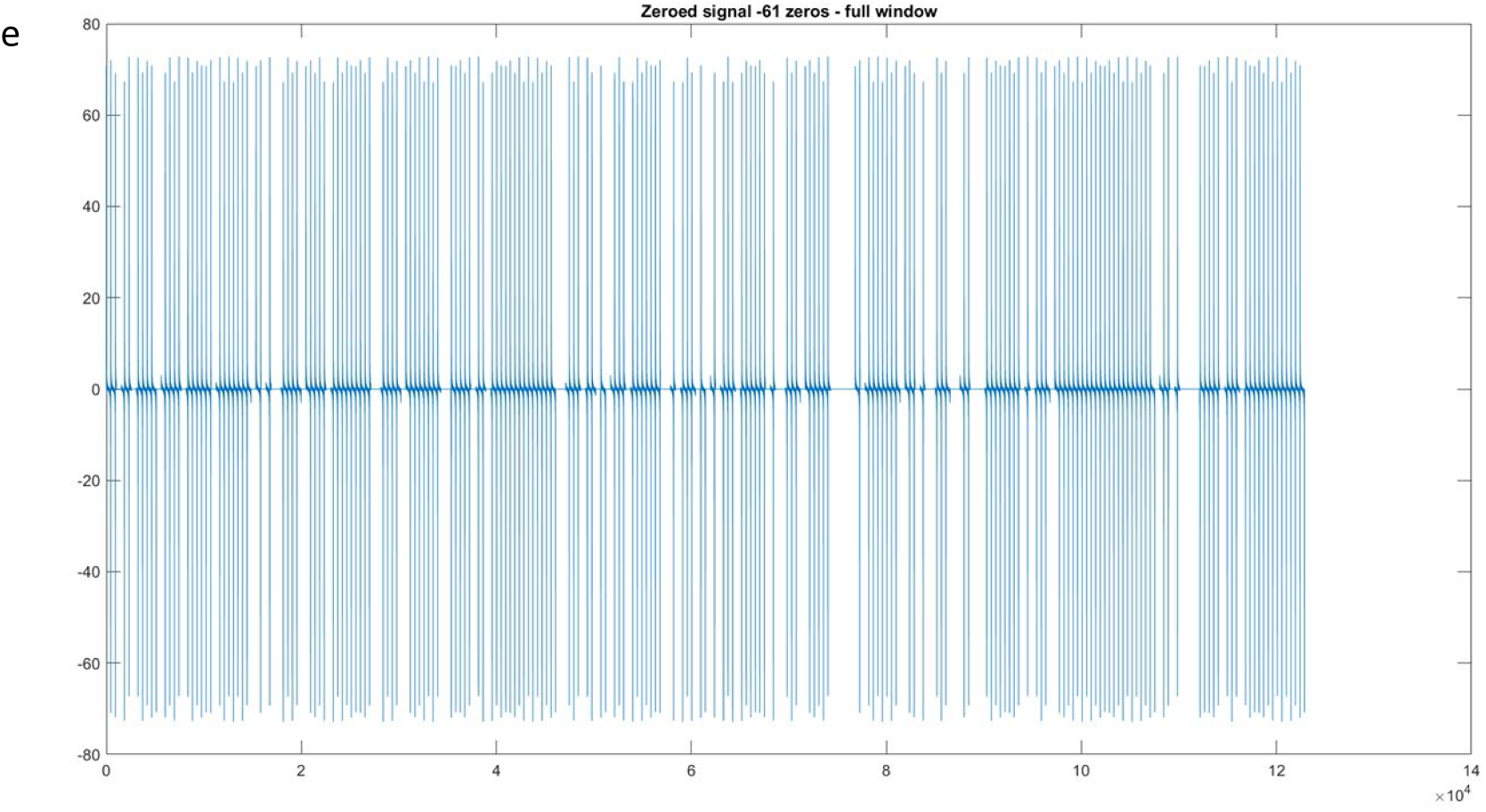

To ensure that this was better than the alternative, white noise was generated which was at least 70% of the amplitude of the sinusoidal waves and randomly added into the original (a) timeseries with up to 25% of the one second segments being replaced. Timeseries with added noise is shown in (f), with a close up (g) and a spectral decomposition (h). There is a lot of distortion to the original signal (a), along with a lot of noise added broadband across the spectra, which suggests that zeroing out noisy segments is worth doing.

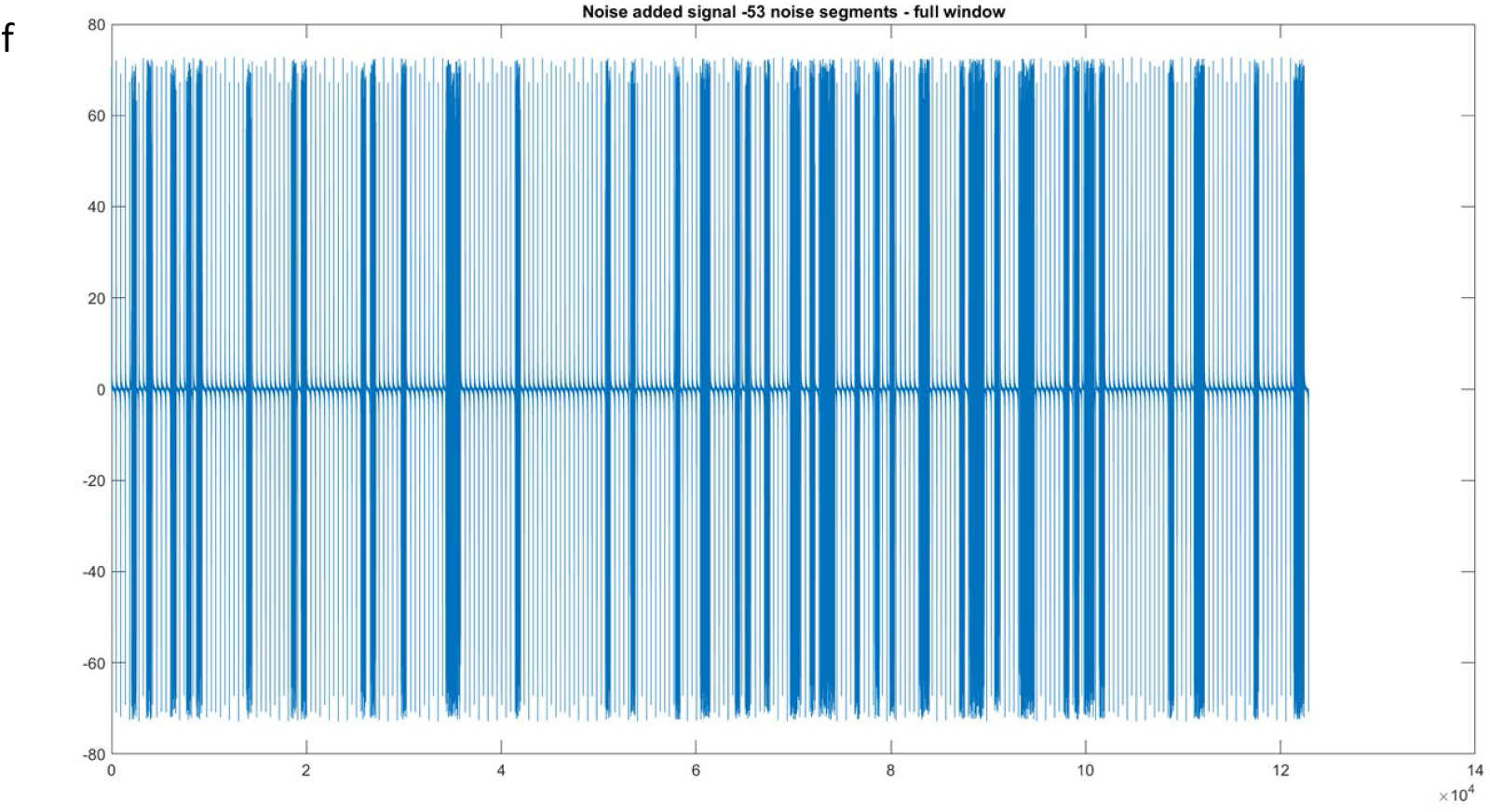
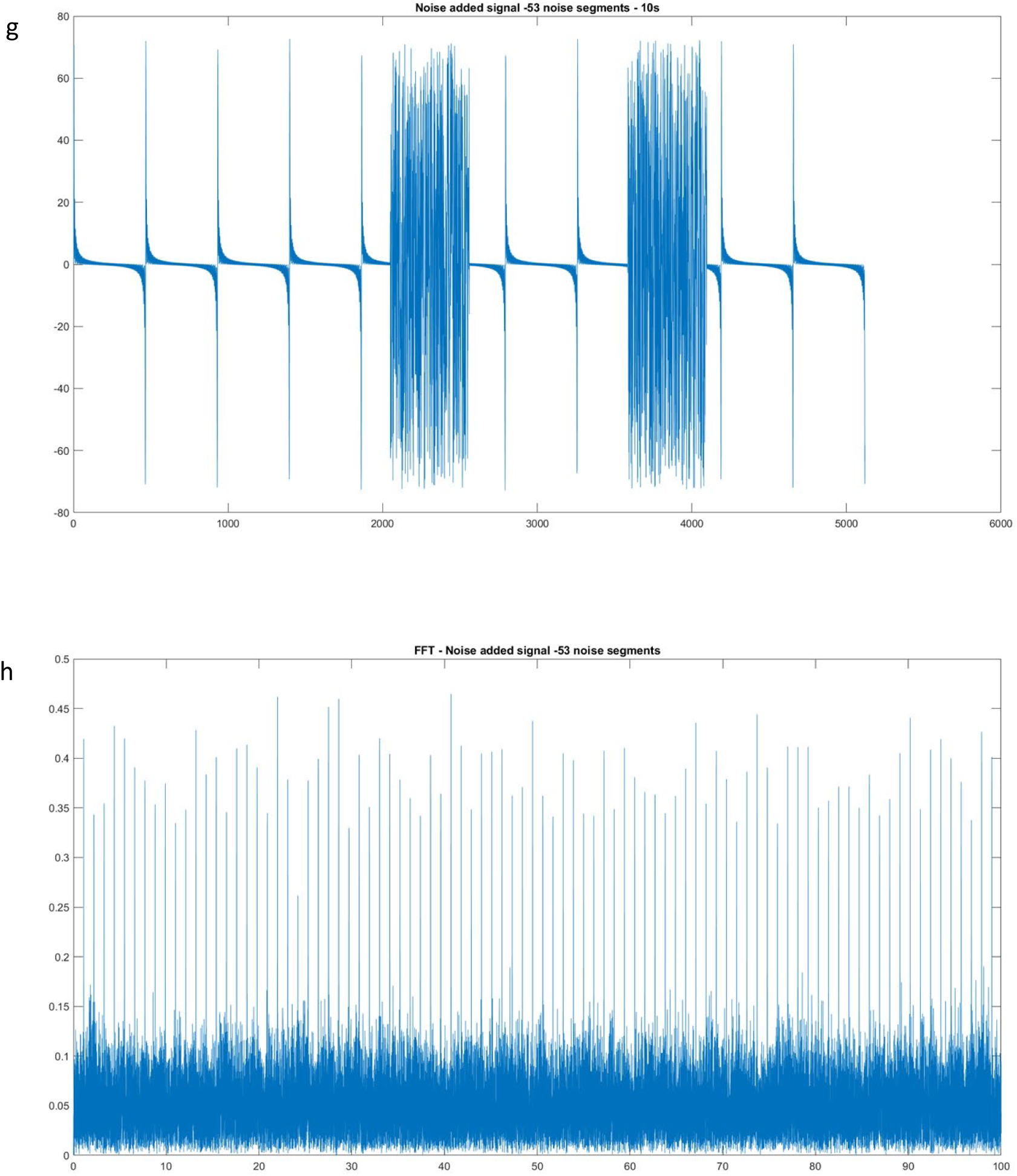

Two alternatives were also tested, first cutting out bad segments and concatenating zeroed out segments to the end of the temporal timeseries before spectral decomposition (i) showed that this greatly distorts the spectral timeseries, likely due to edge effects that are exacerbated.

To combat edge effects a version using a hanning window was trialled timeseries (j), close up (k) and spectral decomposition (l) are shown below. This ended up having the same effect as zeroing out one second segments.

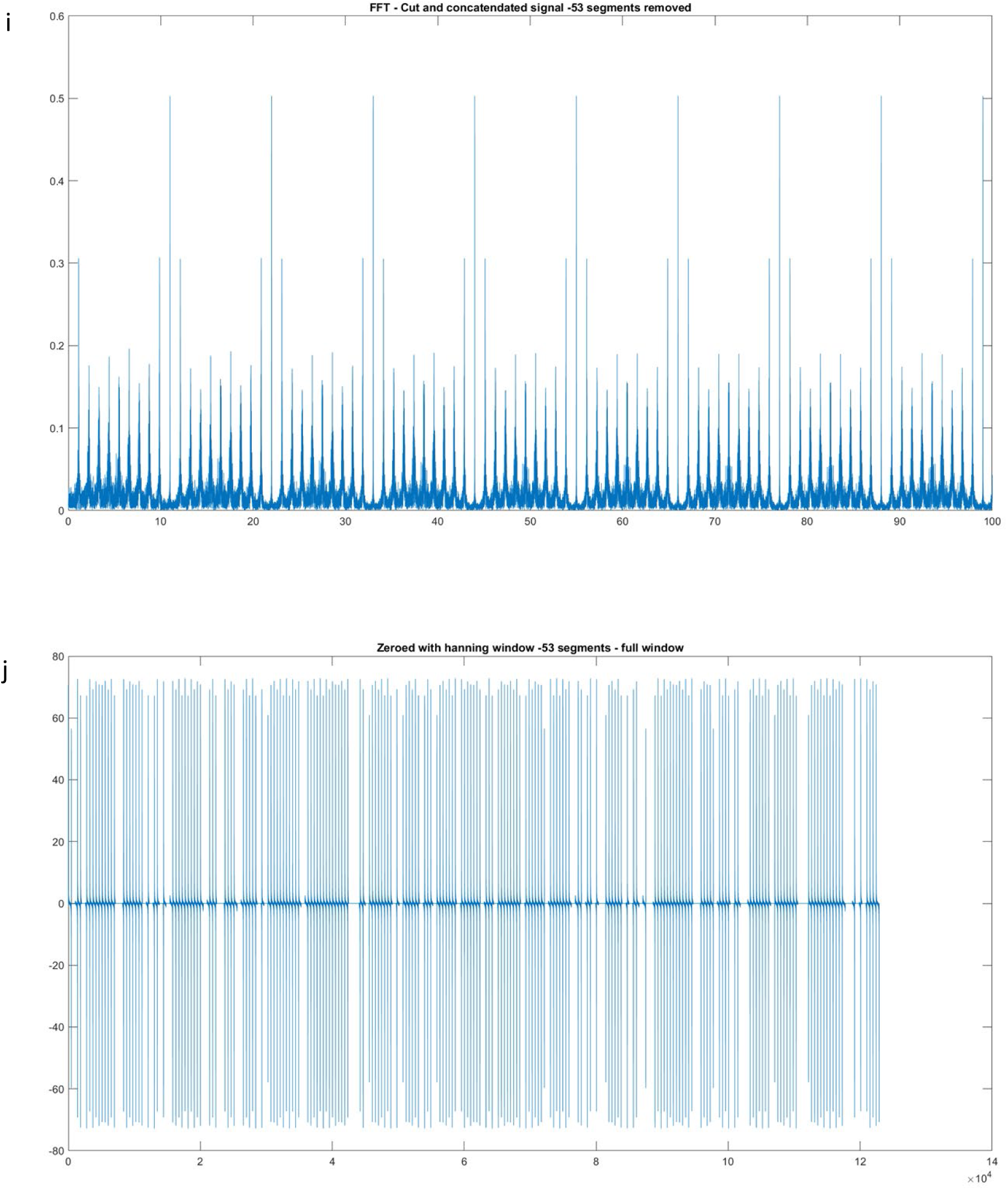
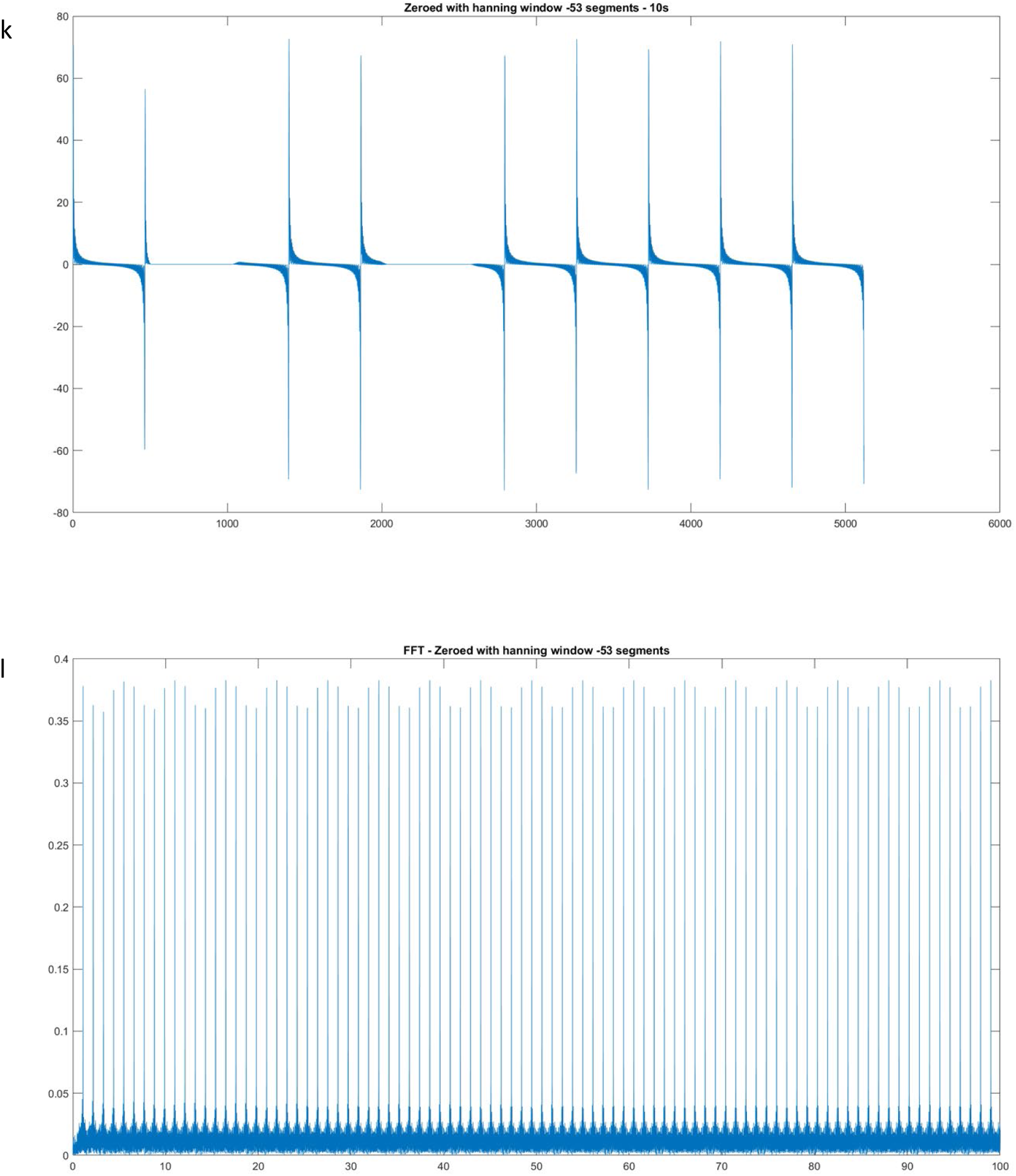

## Data Quality Metrics

To control for data quality issues, all participants were rejected using the criteria in the methods section. The total summed power, percentage of channel interpolation and percentage of sections zeroed due to high noise levels are shown in the table below.

|  | Infant |  |  | Adult |  |  |
| --- | --- | --- | --- | --- | --- | --- |
|  | Summed power (magnitude) | Interpolated channels | Segments removed | Summed power (magnitude) | Interpolated channels | Segments removed |
| **1042** | 118447 | 3% | 0% | 45830 | 0% | 2.4% |
| **1051** | 44484 | 8% | 18.7% | 119151 | 9% | 7.2% |
| **1052** | 75480 | 11% | 0% |  |  |  |
| **1053** | 199324 | 25% | 0% | 38672 | 9% | 11.4% |
| **1055** |  |  |  | 30681 | 5% | 1.5% |
| **1059** |  |  |  | 35778 | 6% | 17.7% |
| **1061** |  |  |  | 25036 | 17% | 6.7% |
| **1062** | 116355 | 25% | 0% |  |  |  |
| **1068** |  |  |  | 24457 | 2% | 24.8% |
| **1079** | 119729 | 6% | 0% |  |  |  |
| **1082** | 81081 | 11% | 3.8% |  |  |  |
| **1084** | 94111 | 11% | 0% |  |  |  |

## References

Adibpour, P., Lebenberg, J., Kabdebon, C., Dehaene-Lambertz, G., & Dubois, J. (2020). Anatomo- functional correlates of auditory development in infancy. Developmental Cognitive Neuroscience, 42, 100752. https://doi.org/10.1016/j.dcn.2019.100752

Adibpour, P., Lebenberg, J., Kabdebon, C., Dehaene-lambertz, G., & Dubois, J. (2020). Developmental Cognitive Neuroscience Anatomo-functional correlates of auditory development in infancy. Developmental Cognitive Neuroscience, 42, 100752. https://doi.org/10.1016/j.dcn.2019.100752

Ahissar, E., Nagarajan, S., Ahissar, M., Protopapas, A., Mahncke, H., Merzenich, M. (2001). Speech comprehension is correlated with temporal response patterns recorded from auditory cortex. PNAS, 98(23), 13367–13372.

Ahlo, K., Sainio, K., Sajaniemi, N., Reinikainen, K. Näätänen, R. (1990). Event-related brain potential of human newborns to pitch change of an acoustic stimulus. Electroencephalography and Clinical Neurophysiology1, *77*, 151–155.

Ahtola, E., Stjerna, S., Tokariev, A., & Vanhatalo, S. (2020). Use of complex visual stimuli allows controlled recruitment of cortical networks in infants. Clinical Neurophysiology, 131(8), 2032– 2040. https://doi.org/10.1016/j.clinph.2020.03.034

Aiken, S. J., & Picton, T. W. (2008). Human Cortical Responses to the Speech Envelope. Ear & Hearing, 29(2), 139–157.

Alaerts, J., Heleen, L., & Hofmann, M. Wouters, J. (2009). Cortical auditory steady-state responses to low modulation rates. Internation Journal of Audiology, 48, 582–593. https://doi.org/10.1080/14992020902894558

Alschuler, D. M., Tenke, C. E., Bruder, G. E., & Kayser, J. (2014). Identifying electrode bridging from electrical distance distributions: A survey of publicly-available EEG data using a new method. Clinical Neurophysiology, 125(3), 484–490. https://doi.org/10.1016/j.clinph.2013.08.024

Aoyagi, M., Kiren, T., Furuse, H., Fuse, T., Suzuki, Y., Yokota, M., & Koike, Y. (1994). Effects of Aging on Amplitude-modulation Following Response Effects of Aging on Amplitude-modulation Following Response. Acta Oto-Laryngologica, 511, 15–22. https://doi.org/10.3109/00016489409128295

Arnal, L. H., Doelling, K. B., & Poeppel, D. (2015). Delta – Beta Coupled Oscillations Underlie Temporal Prediction Accuracy. September, 3077–3085. https://doi.org/10.1093/cercor/bhu103

Arnal, L. H., & Giraud, A. (2012). Cortical oscillations and sensory predictions. Trends in Cognitive Sciences, 16(7), 390–398. https://doi.org/10.1016/j.tics.2012.05.003

Aston-Jones, G., & Cohen, J. D. (2005). An integrative theory of locus coeruleus-norepinephrine function: Adaptive gain and optimal performance. Annual Review of Neuroscience, 28, 403–450. https://doi.org/10.1146/annurev.neuro.28.061604.135709

Attaheri, A., Choisdealbha, Á. N., Liberto, G. M. Di, Rocha, S., Brusini, P., Mead, N., Olawole-Scott, H., Boutris, P., Gibbon, S., Williams, I., Grey, C., Flanagan, S., & Goswami, U. (2021). Delta- and theta-band cortical tracking and phase-amplitude coupling to sung speech by infants. BioRxiv, 2020.10.12.329326. https://www.biorxiv.org/content/10.1101/2020.10.12.329326v4%0Ahttps://www.biorxiv.org/content/10.1101/2020.10.12.329326v4.abstract

Attaheri, A., Ní Choisdealbha, Á., Di Liberto, G. M., Rocha, S., Brusini, P., Mead, N., Olawole-scott, H., Boutris, P., Gibbon, S., Williams, I., Grey, C., Flanagan, S., & Goswami, U. (2022). NeuroImage Delta- and theta-band cortical tracking and phase-amplitude coupling to sung speech by infants. NeuroImage, 247(March 2021), 118698. https://doi.org/10.1016/j.neuroimage.2021.118698

Babiloni, F., & Astolfi, L. (2014). Neuroscience and Biobehavioral Reviews Social neuroscience and hyperscanning techniques : Past, present and future. Neuroscience and Biobehavioral Reviews, 44, 76–93. https://doi.org/10.1016/j.neubiorev.2012.07.006

Backer, K. C., Kessler, A. S., Lawyer, L. A., Corina, D. P., & Miller, L. M. (2019). A novel EEG paradigm to simultaneously and rapidly assess the functioning of auditory and visual pathways. Journal of Neurophysiology, 122(4), 1312–1329. https://doi.org/10.1152/jn.00868.2018

Backer, Kristina C; Kessler, A. S. (2019). A Novel EEG Paradigm to Simultaneously and Rapidly Assess the Functioning of Auditory and Visual Pathways. 530.

Barnet, Ann; Ohlrich, Elizabeth; Weiss, Ira; Shanks, B. (1975). AUDITORY EVOKED POTENTIALS DURING SLEEP IN NORMAL CHILDREN FROM TEN DAYS TO THREE YEARS OF AGE. Electroencephalography and Clinical Neurophysiology, 39, 29–41.

Bednar, A., & Lalor, E. C. (2020). Where is the cocktail party? Decoding locations of attended and unattended moving sound sources using EEG. NeuroImage, 205(August 2019), 116283. https://doi.org/10.1016/j.neuroimage.2019.116283

Begum Ali, J., Charman, T., Johnson, M. H., Jones, E. J. H., Agyapong, M., Bazelmans, T., Dafner, L., Ersoy, M., Gliga, T., Goodwin, A., Haartsen, R., Hendry, A., Holman, R., Kalwarowsky, S., Kolesnik, A., Lloyd-Fox, S., Mason, L., Pasco, G., Pickles, A., … Taylor, C. (2020). Early Motor Differences in Infants at Elevated Likelihood of Autism Spectrum Disorder and/or Attention Deficit Hyperactivity Disorder. Journal of Autism and Developmental Disorders, 50(12), 4367– 4384. https://doi.org/10.1007/s10803-020-04489-1

Bernstein-Ratner, N. (1985). Dissociations between Vowel Durations and Formant Frequency Characteristics. Journal of Speech, Language, and Hearing Research, 28(2), 255–264. https://doi.org/10.1044/jshr.2802.255

Bertoncini, J., Nazzi, T., Cabrera, L., & Lorenzi, C. (2011). Six-month-old infants discriminate voicing on the basis of temporal envelope cues (L). The Journal of the Acoustical Society of America, 129(5), 2761–2764. https://doi.org/10.1121/1.3571424

Besle, J., Schevon, C. A., Mehta, A. D., Lakatos, P., Goodman, R. R., Mckhann, G. M., Emerson, R. G., & Schroeder, C. E. (2011). Tuning of the Human Neocortex to the Temporal Dynamics of Attended Events. The Journal of Neuroscience, 31(9), 3176–3185. https://doi.org/10.1523/JNEUROSCI.4518-10.2011

Boyer, T. W., Harding, S. M., & Bertenthal, B. I. (2020). The temporal dynamics of infants’ joint attention: Effects of others’ gaze cues and manual actions. Cognition, 197(December). https://doi.org/10.1016/j.cognition.2019.104151

Burgess, A. (2013). On the interpretation of synchronization in EEG hyperscanning studies : a cautionary note. Frontiers in Human Neuroscience, 7(December), 1–17. https://doi.org/10.3389/fnhum.2013.00881

Burgess, A. P. (2012). Towards a Unified Understanding of Event-Related Changes in the EEG: The Firefly Model of Synchronization through Cross-Frequency Phase Modulation. PLoS ONE, 7(9). https://doi.org/10.1371/journal.pone.0045630

Burnham, D., & Dodd, B. (2004). Auditory-visual speech integration by prelinguistic infants: Perception of an emergent consonant in the McGurk effect. Developmental Psychobiology, 45(4), 204–220. https://doi.org/10.1002/dev.20032

Buzsáki, G., & Draguhn, A. (2004). Neuronal Oscillations in Cortical Networks. Science, 304(June), 1926–1929. http://science.sciencemag.org/

Cabrera, L., Calcus, A., Labendzki, P., & Lorenzini, I. (2021). Amplitude Modulation Following Response in Normal-Hearing Adults : Is There a Link With the Ability to Perceive Speech in Noise *?* 2021.

Carral, V., Huotilainen, M., Ruusuvirta, T., Fellman, V., Näätänen, R., & Escera, C. (2005). A kind of auditory ‘ primitive intelligence ’ already present at birth. European Journal of Neuroscience, 21(March), 3201–3204. https://doi.org/10.1111/j.1460-9568.2005.04144.x

Cellier, D., Riddle, J., Petersen, I., & Hwang, K. (2021). Developmental Cognitive Neuroscience The development of theta and alpha neural oscillations from ages 3 to 24 years. Developmental Cognitive Neuroscience, 50, 100969. https://doi.org/10.1016/j.dcn.2021.100969

Centre, D., Neuroscience, M., & Duprez, J. (2020). Synchronization between keyboard typing and neural oscillations. 1–33.

Chave, J., Herault, B., & Etc… (1976). Fo r R ev iew On ly Fo r R iew On ly. 6–82.

Chemero, A. (2009). Radical embodied cognitive science. [References]. In (2009).

Choi, D., Batterink, L. J., Black, A. K., Paller, K. A., & Werker, J. F. (2020). Preverbal Infants Discover Statistical Word Patterns at Similar Rates as Adults : Evidence From Neural Entrainment. Association for Psychological Science, 1–13. https://doi.org/10.1177/0956797620933237

Choisdealbha, Á. N., Attaheri, A., Rocha, S., Mead, N., Brusini, P., Gibbon, S., Boutris, P., Grey, C., Flanagan, S., & Goswami, U. (2022). Cortical Oscillations in Pre-verbal Infants Track Rhythmic Speech and Non-speech Stimuli.

Cirelli, L. K., Spinelli, C., Nozaradan, S., & Trainor, L. J. (2016). Measuring Neural Entrainment to Beat and Meter in Infants : Effects of Music Background. Frontiers in Neuroscience, 10(May), 1–11. https://doi.org/10.3389/fnins.2016.00229

Clerkin, E. M., Hart, E., Rehg, J. M., Yu, C., Smith, L. B., & Smith, L. B. (2017). Real-world visual statistics and infants ’ first-learned object names.

Cohen, L. T., Rickards, F. W., & Clark, G. M. (2014). A comparison of steady-state evoked potentials to modulated tones in awake and sleeping humans. Journal of the Acoustical Society of America, 90(5), 2467–2479.

Cohn, J. F., & Tronick, E. Z. (1988). Mother-Infant Face-to-Face Interaction: Influence is Bidirectional and Unrelated to Periodic Cycles in Either Partner’s Behavior. Developmental Psychology, 24(3), 386–392. https://doi.org/10.1037/0012-1649.24.3.386

Colombo, J., & Cheatham, C. L. (2006). The emergence and basis of endogenous attention in infancy and early childhood. In Advances in Child Development and Behavior (Vol. 34). https://doi.org/10.1016/S0065-2407(06)80010-8

Cooper, R. P., & Aslin, R. N. (1994). Developmental Differences in Infant Attention to the Spectral Properties of Infant-directed Speech. Child Development, 65(6), 1663–1677. https://doi.org/10.1111/j.1467-8624.1994.tb00841.x

Cooper, R. P., & Aslin, R. N. (1990). Preference for Infant-directed Speech in the First Month after Birth. Child Development, 61(5), 1584–1595. https://doi.org/10.1111/j.1467-8624.1990.tb02885.x

Crosse, M. J., Butler, J. S., & Lalor, E. C. (2015). Congruent visual speech enhances cortical entrainment to continuous auditory speech in noise-free conditions. Journal of Neuroscience, 35(42), 14195–14204. https://doi.org/10.1523/JNEUROSCI.1829-15.2015

Crosse, M. J., Di Liberto, G. M., Bednar, A., & Lalor, E. C. (2016). The multivariate temporal response function (mTRF) toolbox: A MATLAB toolbox for relating neural signals to continuous stimuli. Frontiers in Human Neuroscience, 10(NOV2016), 1–14. https://doi.org/10.3389/fnhum.2016.00604

Csibra, G., Davis, G., Spratling, M. W., & Johnson, M. H. (2000). Gamma Oscillations and Object Processing in the Infant Brain. 290(November), 5–8.

Cuevas, K., Cannon, E. N., Yoo, K., & Fox, N. A. (2014). The infant EEG mu rhythm : Methodological considerations and best practices. Developmental Review, 34(1), 26–43. https://doi.org/10.1016/j.dr.2013.12.001

Daneshvarfard, F., Moghaddam, H. A., & Dehaene-lambertz, G. (2019). Neurodevelopment and asymmetry of auditory-related responses to repetitive syllabic stimuli in preterm neonates based on frequency-domain analysis. Scientific Reports, 9(October 2018), 10654. https://doi.org/10.1038/s41598-019-47064-0

Daume, J., Wang, P., Maye, A., Zhang, D., & Engel, A. K. (2021). NeuroImage Non-rhythmic temporal prediction involves phase resets of low-frequency delta oscillations. NeuroImage, 224(September 2020), 117376. https://doi.org/10.1016/j.neuroimage.2020.117376

David, O., Harrison, L., & Friston, K. J. (2005). Modelling event-related responses in the brain. NeuroImage, 25(3), 756–770. https://doi.org/10.1016/j.neuroimage.2004.12.030

David, O., Kilner, J. M., & Friston, K. J. (2006). Mechanisms of evoked and induced responses in MEG/EEG. NeuroImage, 31(4), 1580–1591. https://doi.org/10.1016/j.neuroimage.2006.02.034

Debener, S., Minow, F., Emkes, R., Gandras, K., & Vos, M. D. E. (2012). How about taking a low-cost, small, and wireless EEG for a walk*?* 1–5. https://doi.org/10.1111/j.1469-8986.2012.01471.x

Decasper, A. J., & Fifer, W. P. (1980). Of human bonding: Newborns prefer their mothers’ voices. Science, 208(4448), 1174–1176. https://doi.org/10.1126/science.7375928

Delorme, A. (2006). eeg_interp EEGLAB function. The Mathworks, Inc.

Delorme, A., & Makeig, S. (2004). EEGLAB: An open source toolbox for analysis of single-trial EEG dynamics including independent component analysis. Journal of Neuroscience Methods, 134(1), 9–21. https://doi.org/10.1016/j.jneumeth.2003.10.009

Demany, L. (1982). Auditory stream segregation in infancy. Infant Behavior and Development, 5(2–4), 261–276. https://doi.org/10.1016/S0163-6383(82)80036-2

Demos, A. P., Chaffin, R., & Marsh, K. L. (2007). Spontaneous Vs Intentional Entrainment To a Musical Beat. January, 1–4.

Dikker, S., Wan, L., Davidesco, I., Bavel, J. J. Van, Ding, M., Poeppel, D., Dikker, S., Wan, L., Davidesco, I., Kaggen, L., Oostrik, M., Mcclintock, J., & Rowland, J. (2017). Brain-to-Brain Synchrony Tracks Real-World Report Brain-to-Brain Synchrony Tracks Real-World Dynamic Group Interactions in the Classroom. Current Biology, 1–6. https://doi.org/10.1016/j.cub.2017.04.002

Ding, J., Sperling, G., & Srinivasan, R. (2006). Attentional modulation of SSVEP power depends on the network tagged by the flicker frequency. Cerebral Cortex, 16(7), 1016–1029. https://doi.org/10.1093/cercor/bhj044

Ding, N., Melloni, L., Zhang, H., Tian, X., & Poeppel, D. (2015). Cortical tracking of hierarchical linguistic structures in connected speech. Nature Neuroscience, 1–10. https://doi.org/10.1038/nn.4186

Ding, N., & Simon, J. Z. (2012). Neural coding of continuous speech in auditory cortex during monaural and dichotic listening. Journal of Neurophysiology, 107(1), 78–89. https://doi.org/10.1152/jn.00297.2011

Ding, N., & Simon, J. Z. (2013). Power and phase properties of oscillatory neural responses in the presence of background activity. Journal of Computational Neuroscience, 34(2), 337–343. https://doi.org/10.1007/s10827-012-0424-6

Doelling, K., Poeppel, D. (2015). Cortical entrainment to music and its modulation by expertise. PNAS, 6233–6242.

Doelling, K. B., Florencia Assaneo, M., Bevilacqua, D., Pesaran, B., & Poeppel, D. (2019). An oscillator model better predicts cortical entrainment to music. Proceedings of the National Academy of Sciences of the United States of America, 116(20), 10113–10121. https://doi.org/10.1073/pnas.1816414116

Doelling, K. B., Arnal, L. H., Ghitza, O., & Poeppel, D. (2014). NeuroImage Acoustic landmarks drive delta – theta oscillations to enable speech comprehension by facilitating perceptual parsing. NeuroImage, 85, 761–768. https://doi.org/10.1016/j.neuroimage.2013.06.035

Drullman, R., Festen, J. M., & Plomp, R. (1994). Effect of reducing slow temporal modulations on speech reception. Journal of the Acoustical Society of America, 95(5), 2670–2680. https://doi.org/10.1121/1.409836

Efron, B., & Tibshirani, R. (1993). Bootstrp.m MATLAB function. The Mathworks, Inc.

Eisermann, M., Kaminska, A., Moutard, M. L., Soufflet, C., & Plouin, P. (2013). Normal EEG in childhood: From neonates to adolescents. In Neurophysiologie Clinique (Vol. 43, Issue 1, pp. 35– 65). https://doi.org/10.1016/j.neucli.2012.09.091

Fairhurst, M. T., & Dumas, G. (2019). Reciprocity and alignment: quantifying coupling in dynamic interactions. https://doi.org/10.31234/osf.io/nmg4x

Falk, S., & Kello, C. T. (2017). Hierarchical organization in the temporal structure of infant-direct speech and song. Cognition, 163, 80–86. https://doi.org/10.1016/j.cognition.2017.02.017

Fawcett, C., Arslan, M., Falck-Ytter, T., Roeyers, H., & Gredebäck, G. (2017). Human eyes with dilated pupils induce pupillary contagion in infants. Scientific Reports, 7(1), 2–8. https://doi.org/10.1038/s41598-017-08223-3

Feldman, R. (2007). Parent-infant synchrony and the construction of shared timing; physiological precursors, developmental outcomes, and risk conditions. Journal of Child Psychology and Psychiatry2, 48(3/4), 329–354.

Feldman, R. (2006). From biological rhythms to social rhythms: Physiological precursors of mother- infant synchrony. Developmental Psychology, 42(1), 175–188. https://doi.org/10.1037/0012-1649.42.1.175

Feldman, R. (2007). Parent-infant synchrony: Biological foundations and developmental outcomes. Current Directions in Psychological Science, 16(6), 340–345. https://doi.org/10.1111/j.1467-8721.2007.00532.x

Feldman, R., Magori-Cohen, R., Galili, G., Singer, M., & Louzoun, Y. (2011). Mother and infant coordinate heart rhythms through episodes of interaction synchrony. Infant Behavior and Development, 34(4), 569–577. https://doi.org/10.1016/j.infbeh.2011.06.008

Fernald, A., & Mazzie, C. (1991). Prosody and focus in speech to infants and adults.: EBSCOhost. American Psychological Association, Inc., 27(2), 209–221. http://web.b.ebscohost.com.revproxy.brown.edu/ehost/pdfviewer/pdfviewer?sid=a7d41e1d-1263-4ba9-9b94-bbaf722e1f4e%40sessionmgr110&vid=1&hid=107

Fernald, A., & Simon, T. (1984). Expanded intonation contours in mothers’ speech to newborns. Developmental Psychology, 20(1), 104–113. https://doi.org/10.1037/0012-1649.20.1.104

Feven-Parsons, I. M., & Goslin, J. (2018). Electrophysiological study of action-affordance priming between object names. Brain and Language, 184(June), 20–31. https://doi.org/10.1016/j.bandl.2018.06.002

Fló, A., Benjamin, L., Palu, M., & Dehaene-Lambertz, G. (2022). Sleeping neonates track transitional probabilities in speech but only retain the first syllable of words. Scientific Reports, 12(1), 1–13. https://doi.org/10.1038/s41598-022-08411-w

Fuglsang, S. A., Dau, T., & Hjortkjær, J. (2017). Author ’ s Accepted Manuscript. NeuroImage. https://doi.org/10.1016/j.neuroimage.2017.04.026

Galambos, R., Makeig, S., & Talmachoff, P. J. (1981). A 40-Hz auditory potential recorded from the human scalp. Proceedings of the National Academy of Sciences of the United States of America, *78*(4 II), 2643–2647. https://doi.org/10.1073/pnas.78.4.2643

Ghinst, M. Vander, Bourguignon, M., Niesen, M., Wens, V., Hassid, S., Choufani, G., Jousmäki, V., Hari, R., Goldman, S., & De Tiège, X. (2019). Cortical tracking of speech-in-noise develops from childhood to adulthood. Journal of Neuroscience, 39(15), 2938–2950. https://doi.org/10.1523/JNEUROSCI.1732-18.2019

Gibbon, S., Attaheri, A., Ní Choisdealbha, Á., Rocha, S., Brusini, P., Mead, N., Boutris, P., Olawole- Scott, H., Ahmed, H., Flanagan, S., Mandke, K., Keshavarzi, M., & Goswami, U. (2021). Machine learning accurately classifies neural responses to rhythmic speech vs. non-speech from 8-week- old infant EEG. Brain and Language, 220(July 2020), 0–6. https://doi.org/10.1016/j.bandl.2021.104968

Giraud, A. L., & Poeppel, D. (2012). Cortical oscillations and speech processing: Emerging computational principles and operations. Nature Neuroscience, 15(4), 511–517. https://doi.org/10.1038/nn.3063

Glass, L. (2001). Synchronization and rhythmic processes in physiology. Nature, 410(March), 277– 284.

Gliga, T., Farroni, T., & Cascio, C. J. (2019). Social touch: A new vista for developmental cognitive neuroscience? Developmental Cognitive Neuroscience, 35(xxxx), 1–4. https://doi.org/10.1016/j.dcn.2018.05.006

Golumbic, E. M. Z., Ding, N., Bickel, S., Lakatos, P., Schevon, C. A., Mckhann, G. M., Goodman, R. R., Emerson, R., Mehta, A. D., Simon, J. Z., & Poeppel, D. (2012). Article Mechanisms Underlying Selective Neuronal Tracking of Attended Speech at a ‘“ Cocktail Party .”’ Neuron, 77(5), 980– 991. https://doi.org/10.1016/j.neuron.2012.12.037

Gomez-Ramirez, M., Kelly, S. P., Molholm, S., Sehatpour, P., Schwartz, T. H., & Foxe, J. J. (2011). Oscillatory sensory selection mechanisms during intersensory attention to rhythmic auditory and visual inputs: A human electrocorticographic investigation. Journal of Neuroscience, 31(50), 18556–18567. https://doi.org/10.1523/JNEUROSCI.2164-11.2011

Goswami, U., & Leong, V. (2013). Speech rhythm and temporal structure: Converging perspectives? Laboratory Phonology, 4(1), 67–92. https://doi.org/10.1515/lp-2013-0004

Greenberg, S., Carvey, H., Hitchcock, L., & Chang, S. (2003). Temporal properties of spontaneous speech - A syllable-centric perspective. Journal of Phonetics, 31(3–4), 465–485. https://doi.org/10.1016/j.wocn.2003.09.005

Gross, J., Kluger, D. S., Abbasi, O., Chalas, N., Steingräber, N., Daube, C., & Schoffelen, J. M. (2021). Comparison of undirected frequency-domain connectivity measures for cerebro-peripheral analysis. NeuroImage, 245(September), 118660. https://doi.org/10.1016/j.neuroimage.2021.118660

Guo, Y., Bufacchi, R. J., Novembre, G., Kilintari, M., Moayedi, M., Hu, L., & Iannetti, G. D. (2020). Ultralow-frequency neural entrainment to pain. PLoS Biology, 18(4), 1–27. https://doi.org/10.1371/journal.pbio.3000491

Haegens, S., & Zion Golumbic, E. (2018). Rhythmic facilitation of sensory processing: A critical review. Neuroscience and Biobehavioral Reviews, 86(July 2017), 150–165. https://doi.org/10.1016/j.neubiorev.2017.12.002

Hamilton, A. (n.d.). *Hype, hyperscanning and embodied social neuroscience*.

Hari, R., Hämäläinen, M., & Joutsiniemi, S. L. (1989). Neuromagnetic steady-state responses to auditory stimuli. Journal of the Acoustical Society of America, 86(3), 1033–1039. https://doi.org/10.1121/1.398093

Harris, K. D. (2020). Nonsense correlations in neuroscience. BioRxiv, 2020.11.29.402719. https://doi.org/10.1101/2020.11.29.402719

Hasson, U., & Frith, C. D. (2016). Mirroring and beyond: Coupled dynamics as a generalized framework for modelling social interactions. Philosophical Transactions of the Royal Society B: Biological Sciences, 371(1693). https://doi.org/10.1098/rstb.2015.0366

Haufe, S., Meinecke, F., Görgen, K., Dähne, S., Haynes, J. D., Blankertz, B., & Bießmann, F. (2014). On the interpretation of weight vectors of linear models in multivariate neuroimaging. NeuroImage, 87, 96–110. https://doi.org/10.1016/j.neuroimage.2013.10.067

He, W., Donoghue, T., Sowman, P. F., Seymour, R. A., Brock, J., Crain, S., Voytek, B., & Hillebrand, A. (2019). Co-increasing neuronal noise and beta power in the developing brain. BioRxiv, December. https://doi.org/10.1101/839258

Henry, M. J., Herrmann, B., & Grahn, J. A. (2017). What can we learn about beat perception by comparing brain signals and stimulus envelopes? PLoS ONE, 12(2), 1–17. https://doi.org/10.1371/journal.pone.0172454

Henry, M. J., & Obleser, J. (2012). Frequency modulation entrains slow neural oscillations and optimizes human listening behavior. Proceedings of the National Academy of Sciences of the United States of America, 109(49), 20095–20100. https://doi.org/10.1073/pnas.1213390109

Herdman, A. T., Lins, O., Van Roon, P., Stapells, D. R., Scherg, M., & Picton, T. W. (2002). Intracerebral sources of human auditory steady-state responses. Brain Topography, 15(2), 69–86. https://doi.org/10.1023/A:1021470822922

Hessels, R. S. (2020). How does gaze to faces support face-to-face interaction? A review and perspective. Psychonomic Bulletin and Review, 27(5), 856–881. https://doi.org/10.3758/s13423-020-01715-w

Hickey, P., Merseal, H., Patel, A. D., & Race, E. (2020). NeuroImage Memory in time : Neural tracking of low-frequency rhythm dynamically modulates memory formation. NeuroImage, 213(February), 116693. https://doi.org/10.1016/j.neuroimage.2020.116693

Hoehl, S., Fairhurst, M., & Schirmer, A. (2021). Interactional synchrony: Signals, mechanisms and benefits. Social Cognitive and Affective Neuroscience, 16(1–2), 5–18. https://doi.org/10.1093/scan/nsaa024

Hoehl, S., Fairhurst, M., & Schirmer, A. (2020). Interactional synchrony: signals, mechanisms and benefits. Social Cognitive and Affective Neuroscience. https://doi.org/10.1093/scan/nsaa024

Holler, J., Kendrick, K. H., Casillas, M., & Levinson, S. C. (2015). Editorial: Turn-Taking in Human Communicative Interaction. In Frontiers in Psychology (Vol. 6, Issue DEC). https://doi.org/10.3389/fpsyg.2015.01919

Hyafil, A., Fontolan, L., Kabdebon, C., Gutkin, B., & Giraud, A. L. (2015). Speech encoding by coupled cortical theta and gamma oscillations. ELife, 4(MAY), 1–45. https://doi.org/10.7554/eLife.06213

Jaffe, J., Beebe, B., Feldstein, S., Crown, C. L., & Jasnow, M. D. (2001). Rhythms of dialogue in infancy: coordinated timing in development. Monographs of the Society for Research in Child Development, 66(2), i–viii, 1–132. http://www.ncbi.nlm.nih.gov/pubmed/11428150

Jerger, J., Chmiel, R., Frost, J. D., & Coker, N. (1986). Effect of sleep on the auditory steady state evoked potential. Ear and Hearing, 7(4), 240–245. https://doi.org/10.1097/00003446-198608000-00004

Jervis, B. W., Nichols, M. J., Johnson, T. E., Allen, E., & Hudson, N. R. (1983). A Fundamental Investigation of the Composition of Auditory Evoked Potentials. IEEE Transactions on Biomedical Engineering, BME-30(1), 43–50. https://doi.org/10.1109/TBME.1983.325165

Jessen, S., Fiedler, L., Münte, T. F., & Obleser, J. (2019). Quantifying the individual auditory and visual brain response in 7-month-old infants watching a brief cartoon movie. NeuroImage, 202(April), 116060. https://doi.org/10.1016/j.neuroimage.2019.116060

Jessen, S., Fiedler, L., Münte, T. F., & Obleser, J. (2019). NeuroImage Quantifying the individual auditory and visual brain response in 7-month-old infants watching a brief cartoon movie. NeuroImage, 202(July), 116060. https://doi.org/10.1016/j.neuroimage.2019.116060

Jiang, J., Dai, B., Peng, D., Zhu, C., Liu, L., & Lu, C. (2012). Neural Synchronization during Face-to-Face Communication. The Journal of Neuroscience, 32(45), 16064–16069. https://doi.org/10.1523/JNEUROSCI.2926-12.2012

Jones, E. J. H., Venema, K., Lowy, R., Earl, R. K., & Webb, S. J. (2015). Developmental changes in infant brain activity during naturalistic social experiences. Developmental Psychobiology, 57(7), 842– 853. https://doi.org/10.1002/dev.21336

Kabdebon, C., Pena, M., Buiatti, M., & Dehaene-lambertz, G. (2015). Brain & Language Electrophysiological evidence of statistical learning of long-distance dependencies in 8-month- old preterm and full-term infants. Brain and Language, 148, 25–36. https://doi.org/10.1016/j.bandl.2015.03.005

Kabdebon, C., Fló, A., Heering, A. De, & Aslin, R. (2022). NeuroImage The power of rhythms : how steady-state evoked responses reveal early neurocognitive development. NeuroImage, 254(August 2021), 119150. https://doi.org/10.1016/j.neuroimage.2022.119150

Kalashnikova, M., Peter, V., Di Liberto, G. M., Lalor, E. C., & Burnham, D. (2018). Infant-directed speech facilitates seven-month-old infants’ cortical tracking of speech. Scientific Reports, 8(1), 1–8. https://doi.org/10.1038/s41598-018-32150-6

Kalashnikova, M., Peter, V., Liberto, G. M. Di, & Lalor, E. C. (2018). Infant-directed speech facilitates seven-month-old infants ’ cortical tracking of speech. Scientific Reports, April, 1–8. https://doi.org/10.1038/s41598-018-32150-6

Kandylaki, K. D., & Kotz, S. A. (2020). Distinct cortical rhythms in speech and language processing and some more : a commentary on. Language, Cognition and Neuroscience, 0(0), 1–5. https://doi.org/10.1080/23273798.2020.1757729

Kawasaki, M., Yamada, Y., Ushiku, Y., Miyauchi, E., & Yamaguchi, Y. (2013). Inter-brain synchronization during coordination of speech rhythm in human-to-human social interaction. 1– 8. https://doi.org/10.1038/srep01692

Kayhan, E., Nguyen, T., Matthes, D., Langeloh, M., Michel, C., & Jiang, J. (2022). Interpersonal neural synchrony when predicting others ’ actions during a game of rock - paper - scissors. Scientific Reports, 1–11. https://doi.org/10.1038/s41598-022-16956-z

Kayser, C., Wilson, C., Safaai, H., Sakata, S., & Panzeri, S. (2015). Rhythmic auditory cortex activity at multiple timescales shapes stimulus–response gain and background firing. Journal of Neuroscience, 35(20), 7750–7762. https://doi.org/10.1523/JNEUROSCI.0268-15.2015

Kimbrough Oller, D. (2000). The Emergence of the Speech Capability (1st ed.). Taylor & Francis.

Kingsbury, L., Huang, S., Wang, J., Gu, K., Golshani, P., Wu, Y. E., & Hong, W. (2019). Correlated Neural Activity and Encoding of Behavior across Brains of Socially Interacting Animals. Cell, 178(2), 429–446.e16. https://doi.org/10.1016/j.cell.2019.05.022

Kösem, A., Gramfort, A., & Wassenhove, V. Van. (2014). NeuroImage Encoding of event timing in the phase of neural oscillations. NeuroImage, 92, 274–284. https://doi.org/10.1016/j.neuroimage.2014.02.010

Koskinen, M., & Seppä, M. (2014). NeuroImage Uncovering cortical MEG responses to listened audiobook stories. NeuroImage, 100, 263–270. https://doi.org/10.1016/j.neuroimage.2014.06.018

Kothe, C. (2014). Clean_channels EEGLAb plugin. The Mathworks, Inc.

Kothe, C. (2010). clean_windows EEGLAB plugin. The Mathworks, Inc.

Kozulin, P., Almarza, G., Gobius, I., & Richards, L. J. (2016). Prenatal and Postnatal Determinants of Development. Neuromethods, 109, 3–20. https://doi.org/10.1007/978-1-4939-3014-2

Kushnerenko, E., Eponiene, R., Balan, P., Fellman, V., Huotilainen, M., & Näätänen, R. (2002). Maturation of the auditory event-related potentials during the first year of life. NeuroReport, 13(1), 47–51. https://doi.org/10.1097/00001756-200201210-00014

Kuwada, S., Batra, R., & Maher, V. L. (1986). Scalp potentials of normal and hearing-impaired subjects in response to sinusoidally amplitude-modulated tones. Hearing Research, 21(2), 179–192. https://doi.org/10.1016/0378-5955(86)90038-9

Lachaux, J. P., Rodriguez, E., Martinerie, J., & Varela, F. J. (1999). Measuring phase synchrony in brain signals. Human Brain Mapping, 8(4), 194–208. https://doi.org/10.1002/(SICI)1097-0193(1999)8:4<194::AID-HBM4>3.0.CO;2-C

Lachaux, J-P., Rodriguez, E., Martinerie, J., Varela, F. J. (1978). Measureing Phase Synchrony in Brain Signals. Human Brain Mapping, 8, 194–208. https://doi.org/10.1017/S0007680500048066

Lakatos, P., Gross, J., & Thut, G. (2019). A New Unifying Account of the Roles of Neuronal Entrainment. Current Biology, 29(18), R890–R905. https://doi.org/10.1016/j.cub.2019.07.075

Lakatos, P., Karmos, G., Mehta, A. D., Ulbert, I., & Schroeder, C. E. (2008). Entrainment of neuronal oscillations as a mechanism of attentional selection. Science, 320(5872), 110–113. https://doi.org/10.1126/science.1154735

Lakatos, P., Musacchia, G., O’Connel, M. N., Falchier, A. Y., Javitt, D. C., & Schroeder, C. E. (2013). The Spectrotemporal Filter Mechanism of Auditory Selective Attention. Neuron, 77(4), 750–761. https://doi.org/10.1016/j.neuron.2012.11.034

Lankinen, K., Saari, J., Hari, R., & Koskinen, M. (2014). Intersubject consistency of cortical MEG signals during movie viewing. NeuroImage, 92, 217–224. https://doi.org/10.1016/j.neuroimage.2014.02.004

Leong, V., Byrne, E., Clackson, K., Georgieva, S., Lam, S., & Wass, S. (2017). Speaker gaze increases information coupling between infant and adult brains. Proceedings of the National Academy of Sciences of the United States of America, 114(50), 13290–13295. https://doi.org/10.1073/pnas.1702493114

Leong, V., & Goswami, U. (2015). Acoustic-emergent phonology in the amplitude envelope of child- directed speech. PLoS ONE, 10(12), 1–37. https://doi.org/10.1371/journal.pone.0144411

Leong, V., Noreika, V., Clackson, K., Georgieva, S., Brightman, L., Nutbrown, R., Fujita, S., Neale, D., & Wass, S. (2019). Mother-infant interpersonal neural connectivity predicts infants’ social learning. https://doi.org/10.31234/osf.io/gueaq

Leong, V., Stone, M. A., Turner, R. E., & Goswami, U. (2014). A role for amplitude modulation phase relationships in speech rhythm perception. The Journal of the Acoustical Society of America, 136(1), 366–381. https://doi.org/10.1121/1.4883366

Levi, E. C., Folsom, R. C., & Dobie, R. A. (1993). Amplitude-modulation following response (AMFR): Effects of modulation rate, carrier frequency, age, and state. Hearing Research, 68(1), 42–52. https://doi.org/10.1016/0378-5955(93)90063-7

Levitan, R., & Hirschberg, J. (2011). Measuring acoustic-prosodic entrainment with respect to multiple levels and dimensions. Proceedings of the Annual Conference of the International Speech Communication Association, INTERSPEECH, 3081–3084. https://doi.org/10.21437/interspeech.2011-771

Liégeois-Chauvel, C., Lorenzi, C., Trébuchon, A., Régis, J., & Chauvel, P. (2004). Temporal envelope processing in the human left and right auditory cortices. Cerebral Cortex, 14(7), 731–740. https://doi.org/10.1093/cercor/bhh033

Llinas, R. R. (1988). Intrinsic Electrophysiological Properties Central Nervous System Function. Science, 242, 1654–1664.

Lo, C.-W., Tung, T.-Y., Ke, A. H., & Brennan, J. R. (2022). Hierarchy, Not Lexical Regularity, Modulates Low-Frequency Neural Synchrony During Language Comprehension. Neurobiology of Language, 3(4), 538–555. https://doi.org/10.1162/nol_a_00077

Lorenzini, I., Labendzki, P., Hababou, M., & Basire, C. (2022). Human neural processing of auditory temporal modulations during the first year of life. *PsyArXiv*, *(Preprint)*, 1–21.

Luo, H., Wang, Y., Poeppel, D., & Simon, J. Z. (2006). Concurrent encoding of frequency and amplitude modulation in human auditory cortex: MEG evidence. Journal of Neurophysiology, 96(5), 2712–2723. https://doi.org/10.1152/jn.01256.2005

Maitha, C., Goode, J. C., Maulucci, D. P., Lasassmeh, S. M. S., Yu, C., Smith, L. B., & Borjon, J. I. (2020). An open-source, wireless vest for measuring autonomic function in infants.

Makeig, S., Westerfield, M., Jung, T. P., Enghoff, S., Townsend, J., Courchesne, E., & Sejnowski, T. J. (2002). Dynamic brain sources of visual evoked responses. Science, 295(5555), 690–694. https://doi.org/10.1126/science.1066168

Mannix, P., Inwald, D., hathorn, M. Coseloe, K. (1997). Tehrman Entrainment of Heart Rate in the Preterm Infant. Pediatric Research, 42, 282–286.

Marriott Haresign, I., Phillips, E., Whitehorn, M., Noreika, V., Jones, E. J. H., Leong, V., & Wass, S. V. (2021). Automatic classification of ICA components from infant EEG using MARA. Developmental Cognitive Neuroscience, 52(September), 101024. https://doi.org/10.1016/j.dcn.2021.101024

Marshall, P. J., Bar-Haim, Y., & Fox, N. A. (2002). Development of the EEG from 5 months to 4 years of age. Clinical Neurophysiology, 113(8), 1199–1208. https://doi.org/10.1016/S1388-2457(02)00163-3

Masataka, N. (1999). Preference for infant-directed singing in 2-day-old hearing infants of deaf parents. Developmental Psychology, 35(4), 1001–1005. https://doi.org/10.1037/0012-1649.35.4.1001

Maurizi, M., Almadori, G., Paludetti, G., Ottavani, F., Rosignoli, M., Luciano, R. (1990). 40-Hz Steady- State Responses in Newborns and in Children. Audiology, 29, 322–328.

McAdams, S., & Bertoncini, J. (1997). Organization and discrimination of repeating sound sequences by newborn infants. The Journal of the Acoustical Society of America, 102(5), 2945–2953. https://doi.org/10.1121/1.420349

McCulloch, D. L., Orbach, H., & Skarf, B. (1999). Maturation of the pattern-reversal VEP in human infants: A theoretical framework. Vision Research, 39(22), 3673–3680. https://doi.org/10.1016/S0042-6989(99)00091-7

Meyer, L., Sun, Y., & Martin, A. E. (2019). Synchronous, but not entrained : exogenous and endogenous cortical rhythms of speech and language processing. 3798. https://doi.org/10.1080/23273798.2019.1693050

Meyer, L., Sun, Y., Martin, A. E., Meyer, L., Sun, Y., & Entraining, A. E. M. (2020). “ Entraining ” to speech, generating language ? Language, Cognition and Neuroscience, 0(0), 1–11. https://doi.org/10.1080/23273798.2020.1827155

Michel, C. M., & He, B. (2019). EEG source localization. In Handbook of Clinical Neurology (1st ed., Vol. 160). Elsevier B.V. https://doi.org/10.1016/B978-0-444-64032-1.00006-0

Millman, R. E., Prendergast, G., Kitterick, P. T., Woods, W. P., & Green, G. G. R. (2010). Spatiotemporal reconstruction of the auditory steady-state response to frequency modulation using magnetoencephalography. NeuroImage, 49(1), 745–758. https://doi.org/10.1016/j.neuroimage.2009.08.029

Moon, C., Cooper, R. P., & Fifer, W. P. (1993). Two-day-olds prefer their native language. Infant Behavior and Development, 16(4), 495–500. https://doi.org/10.1016/0163-6383(93)80007-U

Mühler, R., Rahne, T., & Verhey, J. L. (2013). Auditory brainstem responses to broad-band chirps: Amplitude growth functions in sedated and anaesthetised infants. International Journal of Pediatric Otorhinolaryngology, 77(1), 49–53. https://doi.org/10.1016/j.ijporl.2012.09.028

Mullen, T. (2012). CleanLine EEGLAB plugin. The Mathworks, Inc.

Murray, L., & Trevarthen, C. (1986). The infant’s role in mother-infant communications. Journal of Child Language, 13(1), 15–29. https://doi.org/10.1017/S0305000900000271

Näätänen, R., Tervaniemi, M., Sussman, E., Paavilainen, P., & Winkler, I. (2001). ‘Primitive intelligence’ in the auditory.pdf. Trends in Neurosciences, 24(5), 283–288.

Narayan, C. R., & McDermott, L. C. (2016). Speech rate and pitch characteristics of infant-directed speech: Longitudinal and cross-linguistic observations. The Journal of the Acoustical Society of America, 139(3), 1272–1281. https://doi.org/10.1121/1.4944634

Neale, D., Georgieva, S., Wass, S., & Leong, V. (2017). Towards a neuroscientific understanding of play : A neuropsychological coding framework for analysing infant-adult play patterns. October. https://doi.org/10.1101/202648

Neuroscience, C. (2019). Effects of maternal singing style on mother–infant arousal and behavior Laura K. Cirelli. 1–27.

Niedźwiecka, A., Ramotowska, S., & Tomalski, P. (2018). Mutual Gaze During Early Mother–Infant Interactions Promotes Attention Control Development. Child Development, 89(6), 2230–2244. https://doi.org/10.1111/cdev.12830

Niepel, D., Krishna, B., Siegel, E. R., Draganova, R., Preissl, H., Govindan, R. B., & Eswaran, H. (2020). A pilot study: Auditory steady-state responses (ASSR) can be measured in human fetuses using fetal magnetoencephalography (fMEG). PLoS ONE, *15*(7 July), 1–19. https://doi.org/10.1371/journal.pone.0235310

Notbohm, A., Kurths, J., & Herrmann, C. S. (2016). Modification of brain oscillations via rhythmic light stimulation provides evidence for entrainment but not for superposition of event-related responses. Frontiers in Human Neuroscience, 10(FEB2016). https://doi.org/10.3389/fnhum.2016.00010

Novak, G. P., Kurtzberg, D., Kreuzer, J. A., & Vaughan, H. G. (1989). Cortical responses to speech sounds and their formants in normal infants: maturational sequence and spatiotemporal analysis. Electroencephalography and Clinical Neurophysiology, 73(4), 295–305. https://doi.org/10.1016/0013-4694(89)90108-9

Nozaradan, S., Keller, P. E., Rossion, B., & Mouraux, A. (2018). EEG Frequency-Tagging and Input– Output Comparison in Rhythm Perception. Brain Topography, 31(2), 153–160. https://doi.org/10.1007/s10548-017-0605-8

Nozaradan, S., Peretz, I., Missal, M., & Mouraux, A. (2011). Tagging the neuronal entrainment to beat and meter. Journal of Neuroscience, 31(28), 10234–10240. https://doi.org/10.1523/JNEUROSCI.0411-11.2011

Nozaradan, S., Peretz, I., & Mouraux, A. (2012). Selective neuronal entrainment to the beat and meter embedded in a musical rhythm. Journal of Neuroscience, 32(49), 17572–17581. https://doi.org/10.1523/JNEUROSCI.3203-12.2012

Olsen, W. O. (1977). Average Speech Levels and Spectra in Various Speaking/Listening Conditions: A Summary of the Pearson, Bennett & Fidell (1977) Report. American Journal of Audiology, 7.

Ostchega, Y. Porter, K., Hughes, J., Dillon, C., Nwankwo, T. (2011). Resting Pulse Rate Reference Data for Children, Adolescents, and Adults: United States, 1999-2008 (pp. 1–16).

Park, H., Ince, R. A. A., Schyns, P. G., Thut, G., Park, H., Ince, R. A. A., Schyns, P. G., Thut, G., & Gross, J. (2015). Frontal Top-Down Signals Increase Coupling of Auditory Low-Frequency Oscillations to Continuous Speech in Human Listeners Report Frontal Top-Down Signals Increase Coupling of Auditory Low-Frequency Oscillations to Continuous Speech in Human Listeners. Current Biology, 25(12), 1649–1653. https://doi.org/10.1016/j.cub.2015.04.049

Paulus, M., Hunnius, S., & Bekkering, H. (2013). Neurocognitive mechanisms underlying social learning in infancy : infants ’ neural processing of the effects of others ’ actions. 774–779. https://doi.org/10.1093/scan/nss065

Peelle, J. E., Gross, J., & Davis, M. H. (2013). Phase-locked responses to speech in human auditory cortex are enhanced during comprehension. Cerebral Cortex, 23(6), 1378–1387. https://doi.org/10.1093/cercor/bhs118

Peelle, J. E., Gross, J., & Davis, M. H. (2013). Phase-Locked Responses to Speech in Human Auditory Cortex are Enhanced During Comprehension. June, 1378–1387. https://doi.org/10.1093/cercor/bhs118

Pérez, A., Carreiras, M., & Duñabeitia, J. A. (2017). Brain-To-brain entrainment: EEG interbrain synchronization while speaking and listening. Scientific Reports, 7(1), 1–12. https://doi.org/10.1038/s41598-017-04464-4

Perone, S., & Gartstein, M. A. (2019). Relations between dynamics of parent-infant interactions and baseline EEG functional connectivity. Infant Behavior and Development, 57(March), 101344. https://doi.org/10.1016/j.infbeh.2019.101344

Pethe, J., Mühler, R., Siewert, K., & Von Specht, H. (2004). Near-threshold recordings of amplitude modulation following responses (AMFR) in children of different ages. International Journal of Audiology, 43(6), 339–345. https://doi.org/10.1080/14992020400050043

Picton, T., Sasha John, M., Dimitrijevic, A., Purcell, D. (2003). Human auditory steady-state responses. International Journal of Audiology, 42, 177–219.

Picton, T. W., Skinner, C. R., Champagne, S. C., & Kellett, A. J. C. (1987). Potentials evoked by the sinusoidal modulation of the amplitude or frequency of a tone. Journal of Acousitic Society of America, 82(1), 165–178.

Poulin-Dubois, Diane; Brosseau-Liard, P. (2017). The Developmental Origins of Selective Social Learning. Physiology & Behavior, 176(3), 139–148. https://doi.org/10.1177/0963721415613962.The

Povel, D., & Essens, P. (1985). Perception of Temporal Patterns. Music Perception, 2(4), 411–440.

Power, A. J., Mead, N., Barnes, L., & Goswami, U. (2012). Neural Entrainment to Rhythmically Presented Auditory, Visual, and Audio-Visual Speech in Children. Frontiers in Psychology, 3(July), 1–13. https://doi.org/10.3389/fpsyg.2012.00216

Rees, A., Green, G. G. R., & Kay, R. H. (1986). Steady-state evoked responses to sinusoidally amplitude-modulated sounds recorded in man. Hearing Research, 23(2), 123–133. https://doi.org/10.1016/0378-5955(86)90009-2

Regan, D. (1989). Human brain electrophysiology: Evoked potentials and evoked magnetic fields in science and medicine.

Reid, V. M. (2007). The directed attention model of infant social cognition. May 2014. https://doi.org/10.1080/17405620601005648

Reid, V. M., Striano, T., & Iacoboni, M. (2011). Developmental Cognitive Neuroscience Neural correlates of dyadic interaction during infancy. Accident Analysis and Prevention, 1(2), 124–130. https://doi.org/10.1016/j.dcn.2011.01.001

Reindl, V., Gerloff, C., Scharke, W., & Konrad, K. (2018). SC. NeuroImage. https://doi.org/10.1016/j.neuroimage.2018.05.060

Rekow, D., Baudouin, J. Y., Durand, K., & Leleu, A. (2022). Smell what you hardly see: Odors assist visual categorization in the human brain. NeuroImage, 255(January), 119181. https://doi.org/10.1016/j.neuroimage.2022.119181

Repp, B. H. (2006). Rate Limits of Sensorimotor Synchronization. 2(2), 163–181.

Retter, T. L., Rossion, B., & Schiltz, C. (2021). Harmonic amplitude summation for frequency-tagging analysis. Journal of Cognitive Neuroscience, 33(11), 2372–2393. https://doi.org/10.1162/jocn_a_01763

Rickards, F. W., Tan, L. E., Cohen, L. T., Wilson, O, J., Drew, J. H., Clark, G. M. (1994). Auditory steady-state evoked potentials in newborns. British Journal of Audiology1, 28, 327–337.

Rimmele, J. M., Morillon, B., Poeppel, D., & Arnal, L. H. (2018). Proactive Sensing of Periodic and Aperiodic Auditory Patterns. In Trends in Cognitive Sciences (Vol. 22, Issue 10, pp. 870–882). Elsevier Ltd. https://doi.org/10.1016/j.tics.2018.08.003

Riquelme, R., Kuwada, S., Filipovic, B., Hartung, K., & Leonard, G. (2006). Optimizing the stimuli to evoke the amplitude modulation following response (AMFR) in neonates. Ear and Hearing, 27(2), 104–119. https://doi.org/10.1097/01.aud.0000201857.99240.24

Robertson, S. S., Watamura, S. E., & Wilbourn, M. P. (2012). Attentional dynamics of infant visual foraging. Proceedings of the National Academy of Sciences of the United States of America, 109(28), 11460–11464. https://doi.org/10.1073/pnas.1203482109

Rojas, D. C., Maharajh, K., Teale, P. D., Kleman, M. R., Benkers, T. L., Carlson, J. P., & Reite, M. L. (2006). Development of the 40 Hz steady state auditory evoked magnetic field from ages 5 to 52. Clinical Neurophysiology, 117(1), 110–117. https://doi.org/10.1016/j.clinph.2005.08.032

Rosen, S. (1992). Temporal information in speech: acoustic, auditory and linguistic aspects. In Philosophical transactions of the Royal Society of London. Series B, Biological sciences (Vol. 336, Issue 1278, pp. 367–373). https://doi.org/10.1098/rstb.1992.0070

Roß, B., Borgmann, C., Draganova, R., Roberts, L. E., & Pantev, C. (2000). A high-precision magnetoencephalographic study of human auditory steady-state responses to amplitude- modulated tones. The Journal of the Acoustical Society of America, 108(2), 679–691. https://doi.org/10.1121/1.429600

Saby, J. N., & Marshall, P. J. (2012). Developmental Neuropsychology The Utility of EEG Band Power Analysis in the Study of Infancy and Early Childhood The Utility of EEG Band Power Analysis in the Study of Infancy and Early Childhood. December, 37–41. https://doi.org/10.1080/87565641.2011.614663

Saffran, J. R., & Kirkham, N. Z. (2018). Infant Statistical Learning. Annual Review of Psychology, 69, 181–203. https://doi.org/10.1146/annurev-psych-122216-011805

Saffran, J. R., Werker, J. F., & Werner, L. A. (2007). The Infant’s Auditory World: Hearing, Speech, and the Beginnings of Language. In Handbook of Child Psychology (pp. 58–108). https://doi.org/10.1002/9780470147658.chpsy0202

Saffran, J. R., Johnson, E. K., Aslin, R. N., & Newport, E. L. (1999). Statistical learning of tone sequences by human infants and adults. Cognition, 70(1), 27–52. https://doi.org/10.1016/S0010-0277(98)00075-4

Sammon, M. P., & Darnall, R. A. (1994). Entrainment of respiration to rocking in premature infants: Coherence analysis. Journal of Applied Physiology, 77(3), 1548–1554. https://doi.org/10.1152/jappl.1994.77.3.1548

Sauseng, P., Klimesch, W., Gruber, W. R., Hanslmayr, S., Freunberger, R., & Doppelmayr, M. (2007). Are event-related potential components generated by phase resetting of brain oscillations? A critical discussion. Neuroscience, 146(4), 1435–1444. https://doi.org/10.1016/j.neuroscience.2007.03.014

Savio, G., Cárdenas, J., Pérez Abalo, M. C., González, A., & Valdés, J. (2001). The low and high frequency auditory steady state responses mature at different rates. Audiology and Neuro- Otology, 6(5), 279–287. https://doi.org/10.1159/000046133

Schaworonkow, N., & Voytek, B. (2021). Longitudinal changes in aperiodic and periodic activity in electrophysiological recordings in the first seven months of life. Developmental Cognitive Neuroscience, 47, 100895. https://doi.org/10.1016/j.dcn.2020.100895

Schroeder, C. E., & Lakatos, P. (2009). Low-frequency neuronal oscillations as instruments of sensory selection. Trends in Neurosciences, 32(1), 9–18. https://doi.org/10.1016/j.tins.2008.09.012

Schroeder, C. E., Lakatos, P., Kajikawa, Y., Partan, S., & Puce, A. (2008). Neuronal oscillations and visual amplification of speech. Trends in Cognitive Sciences, 12(3), 106–113. https://doi.org/10.1016/j.tics.2008.01.002

Service, B., Disclosure, T., Service, B., Act, O., Certificate, E. D. B. S., Dbs, E., Certificate, D. B. S., Certificate, D. B. S., Ambassador, U. E. L. S., If, M., London, E., & Onlinedisclosures, G. B. G. (2019). Disclosure and Barring Service ( DBS ) Checks – Frequently asked questions. February.

Shafer, V. L., Yu, Y. H., & Wagner, M. (2015). Maturation of cortical auditory evoked potentials (CAEPs) to speech recorded from frontocentral and temporal sites: Three months to eight years of age. International Journal of Psychophysiology, 95(2), 77–93. https://doi.org/10.1016/j.ijpsycho.2014.08.1390

Shannon, C. (1948). A Mathematical Theory of Communication. The Bell System Technical Journal, 196(4), 519–520. https://doi.org/10.1016/s0016-0032(23)90506-5

Sheean, et al., 2013. (2008). 基因的改变NIH Public Access. Bone, 23(1), 1–7. https://doi.org/10.1038/jid.2014.371

Singh, N. C., & Theunissen, F. E. (2003). Modulation spectra of natural sounds and ethological theories of auditory processing. The Journal of the Acoustical Society of America, 114(6), 3394–3411. https://doi.org/10.1121/1.1624067

Smith, L. B., & Thelen, E. (2003). Development as a dynamic system. Trends in Cognitive Sciences, 7(8), 343–348. https://doi.org/10.1016/S1364-6613(03)00156-6

Smith, L. B., Jayaraman, S., Clerkin, E., & Yu, C. (2018). The Developing Infant Creates a Curriculum for Statistical Learning. Trends in Cognitive Sciences, 22(4), 325–336. https://doi.org/10.1016/j.tics.2018.02.004

Smith, N. A., & Trainor, L. J. (2008). Infant-directed speech is modulated by infant feedback. Infancy, 13(4), 410–420. https://doi.org/10.1080/15250000802188719

Stapells, D. R., Galambos, R., Costello, J. A., & Makeig, S. (1988). Inconsistency of auditory middle latency and steady-state responses in infants 1 David R. Stapells, Robert Galambos, Jamie A. Costello and Scott Makeig. Electroencephalography and Clinical Neurophysiology, 71, 289–295.

Stefanics, G., Haden, G., Huotilainen, M., Balazs, L., Sziller, I., Beke, A., Fellman, V., Winkler, I. (2007). Auditory temporal grouping in newborn infants. Psychophysiology, 44, 697–702.

Stenberg, G. (2013). Do 12-month-old infants trust a competent adult? Infancy, 18(5), 873–904. https://doi.org/10.1111/infa.12011

Study, A. E. E. G. (2020). brain sciences Dynamic Causal Modelling of the Reduced Habituation to Painful Stimuli in Migraine :

Suanda, S. H., Smith, L. B., & Yu, C. (2017). The Multisensory Nature of Verbal Discourse in Parent – Toddler Interactions The Multisensory Nature of Verbal Discourse in Parent – Toddler. Developmental Neuropsychology, 41(5–8), 324–341. https://doi.org/10.1080/87565641.2016.1256403

Tallon-Baudry, C., Bertrand, O., Delpuech, C., & Pernier, J. (1996). Stimulus specificity of phase-locked and non-phase-locked 40 Hz visual responses in human. Journal of Neuroscience, 16(13), 4240– 4249. https://doi.org/10.1523/jneurosci.16-13-04240.1996

Tamis-LeMonda, C. (2006). Child Psychology: A Handbook of Contemporary Issues (C. Balter, L., Tamis-LeMOnda (ed.); 2nd ed.). Psychology Press, Taylor & Francis Group.

Tang, J.-S.-Y., & Maidment, J.-A. (1996). Prosodic Aspects of Child-Directed Speech in Cantonese. Speech, Hearing and Language, 9(1989), 257–276.

Tass, P., Rosenblum, M. G., Weule, J., Kurths, J., Pikovsky, A., Volkmann, J., Schnitzler, A., & Freund, H. (1998). Detection of n:m Phase Locking from Noisy Data: Application to Magnetoencephalography. Physical Review Letters, 81(15), 3291–3294. papers3://publication/uuid/EE6C3B36-05D2-46A0-A8D0-6E09C3C2BF78

Technische Universtität München, L.-M.-U. M. (2018). 済無No Title No Title. In e-conversion - Proposal for a Cluster of Excellence.

Ten Oever, S., Schroeder, C. E., Poeppel, D., Van Atteveldt, N., Mehta, A. D., Mégevand, P., Groppe, D. M., & Zion-Golumbic, E. (2017). Low-frequency cortical oscillations entrain to subthreshold rhythmic auditory stimuli. Journal of Neuroscience, 37(19), 4903–4912. https://doi.org/10.1523/JNEUROSCI.3658-16.2017

Thiessen, E. D., Hill, E. A., & Saffran, J. R. (2005). Infant-directed speech facilitates word segmentation. Infancy, 7(1), 53–71. https://doi.org/10.1207/s15327078in0701_5

Thut, G., Schyns, P. G., & Gross, J. (2011). Entrainment of perceptually relevant brain oscillations by non-invasive rhythmic stimulation of the human brain. In Frontiers in Psychology (Vol. 2, Issue JUL, pp. 1–10). https://doi.org/10.3389/fpsyg.2011.00170

Tierney, A., & Kraus, N. (n.d.). nc or re ct ed Pr oo f nc or re ed Pr oo f. https://doi.org/10.1162/jocn

Uhlhaas, P. J., Roux, F., Rodriguez, E., Rotarska-Jagiela, A., & Singer, W. (2010). Neural synchrony and the development of cortical networks. In Trends in Cognitive Sciences (Vol. 14, Issue 2, pp. 72– 80). https://doi.org/10.1016/j.tics.2009.12.002

Vanrullen, R. (2016). Perceptual Cycles. Trends in Cognitive Sciences, 20(10), 723–735. https://doi.org/10.1016/j.tics.2016.07.006

Varlet, M., Williams, R., & Keller, P. E. (2020). Effects of pitch and tempo of auditory rhythms on spontaneous movement entrainment and stabilisation. Psychological Research, 84(3), 568–584. https://doi.org/10.1007/s00426-018-1074-8

Vettori, S., Dzhelyova, M., Van der Donck, S., Jacques, C., Steyaert, J., Rossion, B., & Boets, B. (2020). Frequency-Tagging Electroencephalography of Superimposed Social and Non-Social Visual Stimulation Streams Reveals Reduced Saliency of Faces in Autism Spectrum Disorder. Frontiers in Psychiatry, 11(April), 1–12. https://doi.org/10.3389/fpsyt.2020.00332

Vettori, S., Dzhelyova, M., Van der Donck, S., Jacques, C., Steyaert, J., Rossion, B., & Boets, B. (2019). Reduced neural sensitivity to rapid individual face discrimination in autism spectrum disorder. NeuroImage: Clinical, 21(November 2018), 101613. https://doi.org/10.1016/j.nicl.2018.101613

Von Stein, A., Chiang, C., & König, P. (2000). Top-down processing mediated by interareal synchronization. Proceedings of the National Academy of Sciences of the United States of America, 97(26), 14748–14753. https://doi.org/10.1073/pnas.97.26.14748

Wallaert, N., Moore, B. C. J., & Lorenzi, C. (2016). Comparing the effects of age on amplitude modulation and frequency modulation detection. The Journal of the Acoustical Society of America, 139(6), 3088–3096. https://doi.org/10.1121/1.4953019

Wang, Y., Ding, N., Ahmar, N., Xiang, J., Poeppel, D., Simon, J. (2011). Sensitivity to temporal modulation rate and spectral bandwidth in the human auditory system: MEG evidence. Journal of Neurophysiology, 107, 2033–2041.

Wang, Y., Lu, L., Zou, G., Zheng, L., Qin, L., Zou, Q., & Gao, J. H. (2022). Disrupted neural tracking of sound localization during non-rapid eye movement sleep. NeuroImage, 260(January), 119490. https://doi.org/10.1016/j.neuroimage.2022.119490

Ward, L. M. (2003). Synchronous neural oscillations and cognitive processes. In Trends in Cognitive Sciences (Vol. 7, Issue 12, pp. 553–559). https://doi.org/10.1016/j.tics.2003.10.012

Warreyn, P., Ruysschaert, L., Wiersema, J. R., Handl, A., Pattyn, G., & Roeyers, H. (2013). Infants ’ mu suppression during the observation of real and mimicked goal-directed actions. 2, 173–185. https://doi.org/10.1111/desc.12014

Wass, S. V., Perapoch Amadó, M., & Ives, J. (2022). Oscillatory entrainment to our early social or physical environment and the emergence of volitional control. Developmental Cognitive Neuroscience, 54(November 2021). https://doi.org/10.1016/j.dcn.2022.101102

Wass, S. V., Clackson, K., Georgieva, S. D., Brightman, L., Nutbrown, R., & Leong, V. (2018). Infants’ visual sustained attention is higher during joint play than solo play: is this due to increased endogenous attention control or exogenous stimulus capture? Developmental Science, 21(6). https://doi.org/10.1111/desc.12667

Wass, S. V., Whitehorn, M., Marriott Haresign, I., Phillips, E., & Leong, V. (2020). Interpersonal Neural Entrainment during Early Social Interaction. Trends in Cognitive Sciences, 24(4), 329–342. https://doi.org/10.1016/j.tics.2020.01.006

Wass, S. V., Smith, C. G., Clackson, K., Gibb, C., Eitzenberger, J., & Mirza, F. U. (2019). Parents Mimic and Influence Their Infant’s Autonomic State through Dynamic Affective State Matching. Current Biology, 29(14), 2415–2422.e4. https://doi.org/10.1016/j.cub.2019.06.016

Weinstein, D., Launay, J., Pearce, E., Dunbar, R. I. M., & Stewart, L. (2016). Singing and social bonding: Changes in connectivity and pain threshold as a function of group size. Evolution and Human Behavior, 37(2), 152–158. https://doi.org/10.1016/j.evolhumbehav.2015.10.002

Wen, X., Mo, J., & Ding, M. (2012). NeuroImage Exploring resting-state functional connectivity with total interdependence. NeuroImage, 60(2), 1587–1595. https://doi.org/10.1016/j.neuroimage.2012.01.079

Widmann, A. (2008). Pop_eegfiltnew EEGLAB plugin. The Mathworks, Inc.

Winkler, I., Kushnerenko, E., Hovárth, J., Čeponienė, R. Fellman, V., Huotilainen, M., Näätänen, R., S. S. (2003). Newborn infants can organize the auditory world. PNAS, 100(20), 11812–11815.

Winkler, I., Háden, G. P., Ladinig, O., Sziller, I., & Honing, H. (2009). Newborn infants detect the beat in music. Proceedings of the National Academy of Sciences of the United States of America, 106(7), 2468–2471. https://doi.org/10.1073/pnas.0809035106

Wunderlich, J. L., & Cone-Wesson, B. K. (2006). Maturation of CAEP in infants and children: A review. Hearing Research, 212(1–2), 212–223. https://doi.org/10.1016/j.heares.2005.11.008

Xie, W., Mallin, B. M., & Richards, J. E. (2018). Development of infant sustained attention and its relation to EEG oscillations: an EEG and cortical source analysis study. Developmental Science, 21(3). https://doi.org/10.1111/desc.12562

Xie, W., & Richards, J. E. (2017). The Relation between Infant Covert Orienting, Sustained Attention and Brain Activity. Brain Topography, 30(2), 198–219. https://doi.org/10.1007/s10548-016-0505-3

Yu, C., & Smith, L. B. (2017). Multiple Sensory-Motor Pathways Lead to Coordinated Visual Attention. 41, 5–31. https://doi.org/10.1111/cogs.12366

Yu, C., & Smith, L. B. (2013). Joint attention without gaze following: Human infants and their parents coordinate visual attention to objects through eye-hand coordination. PLoS ONE, 8(11). https://doi.org/10.1371/journal.pone.0079659

Zhou, H., Melloni, L., Poeppel, D., & Ding, N. (2016). Interpretations of frequency domain analyses of neural entrainment: Periodicity, fundamental frequency, and harmonics. Frontiers in Human Neuroscience, 10(June), 1–8. https://doi.org/10.3389/fnhum.2016.00274

Zoefel, B., & Heil, P. (2013). Detection of near-threshold sounds is independent of eeg phase in common frequency bands. Frontiers in Psychology, 4(MAY), 1–17. https://doi.org/10.3389/fpsyg.2013.00262

Zoefel, B., ten Oever, S., & Sack, A. T. (2018). The involvement of endogenous neural oscillations in the processing of rhythmic input: More than a regular repetition of evoked neural responses. In Frontiers in Neuroscience (Vol. 12, Issue MAR, pp. 1–13). https://doi.org/10.3389/fnins.2018.00095

Zoefel, B., & Vanrullen, R. (2015). The Role of High-Level Processes for Oscillatory Phase Entrainment to Speech Sound. 9(December), 1–12. https://doi.org/10.3389/fnhum.2015.00651

